# Dynamic transcriptional and epigenetic changes define postnatal tendon growth

**DOI:** 10.1101/2024.09.24.614830

**Authors:** Heather L. Dingwall, Mor Grinstein, Terence D. Capellini, Jenna L. Galloway

## Abstract

Tendons are dynamic structures that efficiently transmit forces and enable movement. From birth, tendons undergo dramatic changes from a principally cellular tissue to a hypocellular one characterized by a dense and highly ordered extracellular matrix. During this time, tendon cells change morphology from rounded to stellate in appearance and their proliferative rates decline. There is also significant expansion and maturation of the extracellular matrix (ECM) as tendons grow in length and diameter and alter their biomechanical properties to sustain increased physical activities. Surprisingly, for such an important stage of tendon maturation, we understand very little about the transcriptional and epigenetic regulators that direct these processes. Here, we present a roadmap of genes that are differentially regulated during the early neonatal and postnatal time period. We find differentially expressed genes fall into specific transcriptional modules, representing expression increases, decreases, or gene sets undergoing dynamic changes over postnatal time. By pairing our transcriptomic data with epigenetic data, we were able to perform an integrative analysis of the datasets and further define modules with highly correlated changes in gene expression and chromatin accessibility. From this analysis, several new pathways emerge. Among them, we focus on Yap1, a transcriptional co-activator of the Hippo signaling pathway. We observe accessible regions near to differentially expressed genes, containing motifs for TEAD, the transcription factor that binds Yap to regulate transcription. Conditional loss of *Yap1* at postnatal stages results in altered expression of *Col1a1* and disrupted matrix organization and density, suggesting that Yap is important for refining tendon ECM maturation. Together, our analyses identify a regulator of matrix maturation and provides a rich dataset with which to interrogate transcriptional networks and pathways during this poorly understood time in tendon growth.

## Introduction

Tendons are crucial components of the musculoskeletal system that transmit muscular forces to bones, thus enabling movement. Their highly ordered matrix is mainly composed of type I collagen fibrils and is critical to their ability to efficiently transfer force. Adult tendons are prone to injury, from acute rupture to chronic degeneration, yet following injury, tendons undergo a slow, scar-mediated healing process, resulting in imperfect tissue repair (1). Despite advances in surgical intervention for tendon injuries, the repaired tissue remains mechanically compromised due to this form of healing, yielding a high failure rate after surgery (2, 3). A greater understanding of tendon formation and growth would be instrumental towards eventually developing regenerative based therapies for treating their injury and disease.

Previous work has identified several transcription factors (TF) and signaling pathways that are vital to proper tendon formation. Scleraxis (Scx) is a basic helix-loop-helix TF that has been identified as a marker of tendon cell fate (4). Along with Scx, Mohawk (Mkx) and the Early Growth Response-like (Egr) TFs Egr1 and Egr2 are also important for tendon matrix maturation (5, 6). All four of these TFs can regulate the production of major collagens (Col1a1, Col1a2, Col3a1) and tenomodulin (Tnmd), a transmembrane glycoprotein found on differentiating tendon cells (5–10). TGF-/3 signaling also plays an important role in embryonic tendon formation through the regulation of collagen and other ECM proteins (11–13) and in maintaining *Scx* expression (13–17).

Postnatal tendon growth is characterized by structural and compositional changes to the extracellular matrix (ECM) that produce a highly organized, matrix-rich mature tissue (18–20). Both linear and lateral growth in the tendon continue beyond the highly proliferative postnatal phase indicating a role for ECM expansion during postnatal tendon growth (19–21). There is also detectable cell proliferation in early postnatal days (P) 0 to 14. However, proliferation decreases significantly by P21 and continues at an extremely low rate from 1 month to 1-2 years of life (22).

Interestingly, this transition in cell proliferation and matrix maturation correlates with the timing of a shift in regenerative potential in mammalian tendons. For example, mice, in their first 1-2 weeks of postnatal life, have considerably improved healing potential when compared with 3-week-old (23) and 4–5-month-old mice (24). Tendon healing occurs without the formation of a lasting fibrotic scar in both fetal (2, 25) and early postnatal tendon (18, 24). This regenerative ability is retained even when the immature tendon is wounded after transplantation into the adult environment (2, 25) suggesting the regenerative potential is intrinsic to the tendon. However, this regenerative ability is restricted to fetal and early postnatal stages and declines significantly with maturation (2, 24). A similar decline in regenerative ability has been observed in other organs, such as the heart. Cardiac muscle can regenerate after injury prior to 7 days of neonatal development, but no regeneration is observed at later stages (26); this transition coincides with the developmental stage at which the cardiomyocytes lose their ability to proliferate (27). As in the heart, this drop in tendon cell proliferative activity appears to correlate with the transition from regenerative to reparative healing in mouse tendons around 3 weeks of age (23). Thus, the transition in cell cycle activity and regenerative potential may also correlate with the timing at which tendon cells mature and predominantly function in matrix secretion and maturation.

Although several studies have described postnatal changes to the tendon ECM (18, 20, 22, 28), little work has examined the molecular changes taking place in postnatal tendon cells. The identification of gene regulatory programs that are specifically controlled during this process would be of great significance towards understanding the mechanisms by which tendon cells regulate tendon tissue growth and maturation. Some transcriptomic studies have focused on the active transcriptional programs in embryonic tendon cells (29, 30) whereas others have examined gene expression differences in isolated postnatal tendon progenitor cells after expansion under *in vitro* culture conditions (31). In addition, adult studies have examined differences in mutant mouse tendons (32) and genes involved in aging (33). More recently, single cell RNA-sequencing has revealed tendon cell heterogeneity at multiple stages (17, 34–36), albeit such methods capture a small portion of the tendon cell transcriptome. Additionally, as tendon cell growth and healing potential is likely impacted by non-coding regulation, it will be important to identify *cis*-regulatory regions potentially involved in regulating this transition.

In this study, we use an integrative genomics approach to characterize tendon cells as they transition from highly proliferative to relatively quiescent stages. Specifically, we employ RNA sequencing (RNA-seq) and Assay for Transposase Accessible Chromatin and sequencing (ATAC-seq) to identify expressed genes and open chromatin regions, respectively, in mouse tendon cells as they mature from neonatal stages (P0) to early adulthood (P35). Changes in chromatin accessibility throughout tendon growth can help identify temporally specific putative *cis*-regulatory elements involved in the coordination of the transcriptome dynamics involved in the shift in cell proliferative potential. By integrating these transcriptomic and epigenomic signatures, we are able to identify key genes and signaling pathways that mediate the observed shift in tendon cell proliferative potential. As an example of our ability to identify functionally relevant pathways, we identified the Hippo signaling pathway to be differentially regulated during P0-P35 and tested its role in tendon growth. *Yap1*, a transcriptional co-activator in the Hippo signaling pathway, was specifically deleted at postnatal stages using *Scx-CreERt2*, and collagen expression, organization, and cell number were affected. Together, this research applies innovative techniques to the study of a poorly understood process, providing a rich resource of new candidate pathways to investigate in the context of tendon growth, maturation, and injury repair.

## Results

### Transcriptome dynamics during postnatal tendon growth and development

To specifically focus on postnatal tendon stages when there is a transition from a proliferation-based growth program to one driven by ECM and a shift in regenerative potential, we chose to examine gene expression changes that occur during these time periods. We performed RNA-seq on mRNA isolated from whole distal limb tendons, which were collected weekly at six consecutive time points during the early postnatal period (P0 to P35). A Principal Components Analysis (PCA) on normalized gene counts shows separation of P0, P7, and P14 samples along PC1, while samples from mice at P21 and older do not separate clearly from one another (Fig. 1A). It is also notable that P35 has higher within-group variability than the other time points, which could at least partially explain the inability of this time point to separate from P28. Importantly, PCA shows that sequencing pool does not significantly influence sample clustering (Supp. Fig. 1A) indicating that the group in which a sample was sequenced (sequencing pool) accounts for negligible, if any, sample variance. A likelihood ratio test on gene counts using the DESeq2 framework (37) found that approximately 22% of detected genes were differentially expressed (DE) between at least two time points in the 5-week time series (p adj. < 0.05).

**Figure 1:**
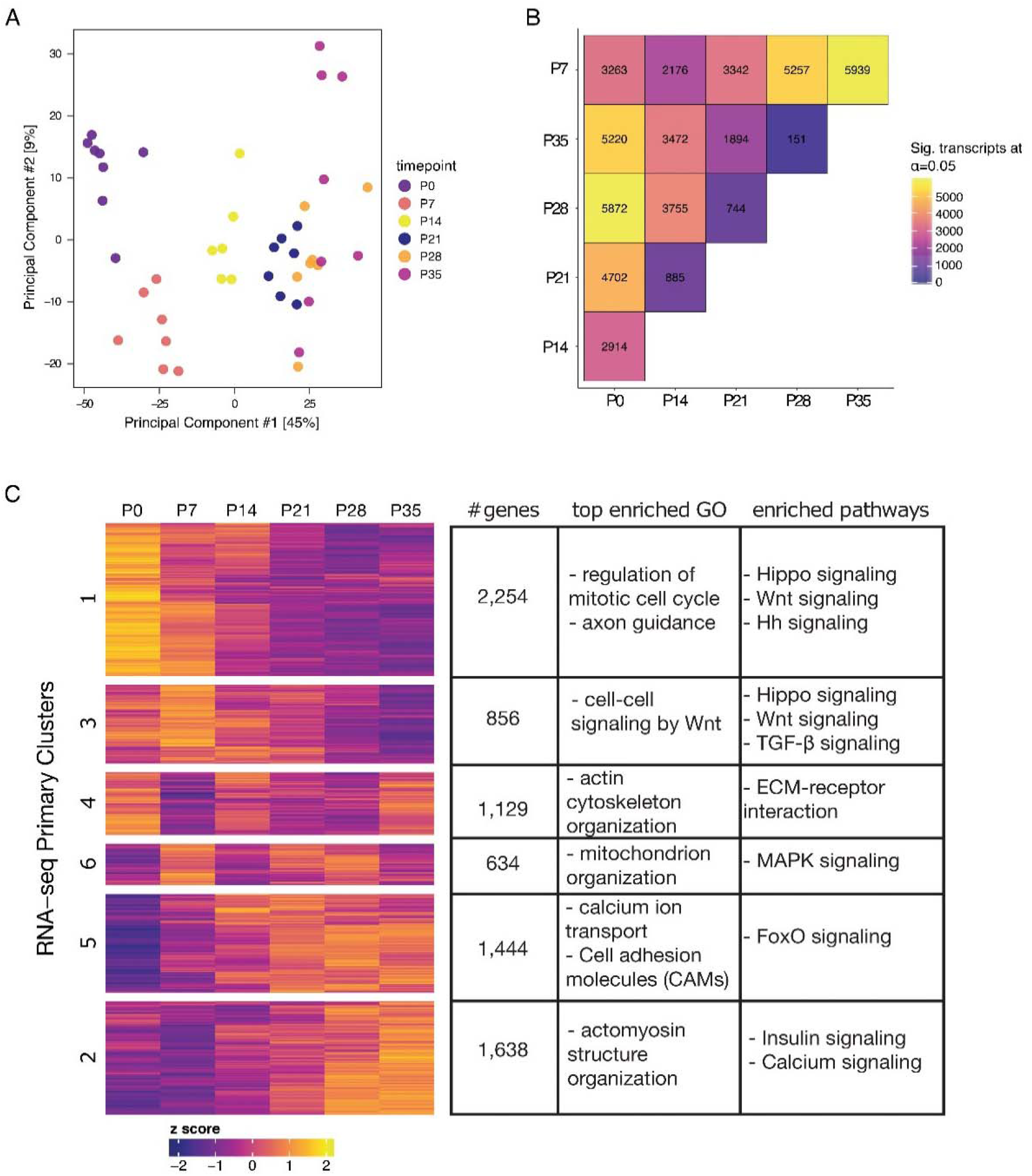
RNA-seq identifies modules of differentially co-expressed genes during postnatal tendon growth. A) PCA of normalized counts from bulk RNA-seq shows separation of timepoints along PC1. Earlier timepoints (P0, P7, and P14) are more distinctly clustered than later timepoints (P21, P28, P35). B) Results of Wald test between all pairs of timepoints (p adj. < 0.05) show more DE genes between pairs of early timepoints compared to pairs of later timepoints. C) Clustered heatmap of genes determined to be DE at any point during the time series (P0 to P35) by likelihood ratio test (p adj. < 0.05); gene clusters were determined using the PAM algorithm. Z-scores were calculated from log2(x+1)-transformed, quantile normalized counts per million reads (CPM). The adjacent table indicates the number of genes occupying each cluster and a selection of the top enrichments from GO and pathway analyses (p adj. < 0.05).

This approach, in which two user-defined models of gene expression are assessed for their goodness of fit, allowed us to test the effects of the time component of the model on gene expression independent of RIN (see Methods), as well as investigate differences in expression among all time points simultaneously. Because our previous work on tendon cell proliferation dynamics (22) points to the second to third postnatal week as an important transitional period during postnatal tendon development, we first investigated the specific pairwise differences in transcriptome-wide gene expression from P0 to P35 and found that many genes are differentially expressed between these three time points (Fig. 1B; Supp Fig. 1C). Given assumptions in the tendon literature that mature tendon cells exhibit relatively low metabolic activity (38), we were quite surprised by the large number of genes (2,508) that were up-regulated from P0 to P35. Comparing the number of differentially expressed genes between each pair of time points shows few genes that are differentially expressed between sequential time points at older stages (i.e., after P14) compared to younger stages (Fig. 1B). This suggests that the first two postnatal weeks are a period of rapid, dynamic transcriptomic change, which slows after P21, mirroring known tendon cell proliferation dynamics (22).

### Expression of known genes involved in tendon development

Next, we defined a set of “tendon” and ECM genes (i.e., genes that code for matrix proteins) based on the current literature to investigate their expression within this data set. Because so few genes are known to be directly involved in tendon development, this set is small enough to examine the expression of each gene individually (Supp. Fig. 2). *Scx*, *Mkx*, *Egr1*, and *Egr2* are all expressed postnatally, but none are significantly DE (Supp. Fig. 2A; p adj. > 0.05). *Scx* expression is highly variable among the individuals within a sample; each time point contains distinct high- and low-expressers of *Scx* (Supp. Fig. 2A). While we have observed higher than expected within-sample variance in *Scx* expression before (22), the cause remains unclear.

We also investigated the expression of TGF-/3 ligands and receptors. The TGF-/3 pathway, and TGFβ2/3 or TGFβR2 specifically, are required for normal tendon development (13). TGFβ signaling is also necessary for the contribution of *Scx*-lineage cells to neonatal regeneration (39). We found that *Tgfb1* is differentially upregulated at P7 only (p adj. < 0.05), while *Tgfb2* and *Tgfb3* are not DE at any point (Supp. Fig. 2B). TGF-/3 receptor expression varies: *Tgfbr1* is intermittently DE, *Tgfbr2* is not DE, and *Tgfbr3* is differentially upregulated from P0 to P35 (p adj. < 0.05; Supp. Fig. 2B).

All ECM related genes in this gene set are significantly DE at some point during the time series (Supp. Fig. 2C; p adj. < 0.05). *Decorin* (*Dcn*) and *Fibromodulin* (*Fmod*) expression increases monotonically from P0 to P35; *Col2a1*, *Col14a1*, *Col3a1*, and *Tenomodulin* (*Tnmd*) expression decreases steadily from P0 to P35; and *Col1a1* and *Col1a2* expression peak around P14 after which they are downregulated. Expression of the matrix gene *Biglycan* (*Bgn*) appears to be partitioned into two phases: it is more highly expressed from P0-P14 followed by downregulation by P21 (p adj. < 0.01), after which expression remains low through P35 (Supp. Fig. 2C).

### Clustering analysis and expression module identification

To gain a more detailed understanding of the temporal gene expression changes that occur between P0 and P35, we performed an unsupervised cluster analysis using the partitioning around medoids (PAM) algorithm on normalized gene counts. This allowed us to discover six differential expression modules – cohorts of genes that tended to be co-expressed over the course of the six time points in the study (Fig. 1C). These co-expression modules reveal 3 broad patterns of differential expression within our data: downregulation over time (Clusters 1 and 4); upregulation over time (Clusters 2 and 5); and intermittent expression (Clusters 3 and 6). The key difference between each pair of expression modules that fit a pattern is the exact timing of the expression change(s). For example, many of the genes grouped into cluster 5 are more strongly upregulated about one week earlier (∼P21) than those in cluster 2 (∼P28). Gene ontology (GO) enrichment analyses on each expression module suggest that biological processes involved in cell proliferation and differentiation dominate the earlier time points, while cell communication and cytoskeleton organization become more important later during postnatal development (Fig. 1C).

Cluster 1, which captures downregulated genes over time, is highly enriched for GO terms related to mitosis, cell cycle, and DNA replication (p. adj < 2 x 10^−18^, q < 5 x 10^−16^). Additionally, a pathway analysis on Cluster 1 found an enrichment of genes involved in the Wnt and Hippo signaling pathways, both of which are known to be involved in regulating cell proliferation in multiple tissues. Like Cluster 1, the genes belonging to Cluster 4 are also most highly expressed at earlier time points. However, most of the genes in this module exhibit peak expression around P7 or P14 instead of P0. Interestingly, enrichment analyses found that the genes comprising Cluster 4 are highly enriched for processes related to the regulation of the actin cytoskeleton, ECM organization, and collagen biosynthesis, as well as small GTPase-mediated signal transduction.

Both modules (Clusters 2 and 5) are characterized by a pattern of steadily increasing expression from P0 to P35 and show enrichment for biological processes involved in calcium ion transport, regulation, and signaling. Clusters 2 and 5 are also enriched for muscle-related GO terms, although they differ in their specifics. Cluster 2 genes, which are upregulated at ∼P28, are enriched for muscle developmental and differentiation processes, as well as actomyosin structure organization. Meanwhile, Cluster 5, which contains genes that are upregulated earlier, is enriched for GO terms related to muscle contraction and structure, in addition to processes involved in calcium ion transport (*S100a1*).

Cluster 3, which contains genes that are expressed intermittently throughout the five-week time series and highest at P0, is also enriched for genes involved in Wnt signaling, as well as Smoothened (Smo) signaling, a key component of the Hedgehog (Hh) signaling pathway. This expression module is also enriched for muscle cell proliferation and differentiation. Multiple GO terms associated with the mitochondria are enriched in the gene set comprising Cluster 6, which is discontinuously upregulated, first at P7 and then from P21 to P28. We also found that Clusters 5 and 6 are enriched for GO terms associated with various metabolic processes and other homeostatic functions, suggesting that the middle of this time series represents the beginning of a shift from growth to homeostasis and a change in cell metabolism.

### Expression of “muscle” genes in the growing tendon

Surprisingly, half of the co-expression modules were enriched for muscle-related GO terms and functions (Fig. 1C). Supp. Fig. 3A illustrates the specific expression patterns of each category of muscle-associated genes identified in Clusters 1, 2, and 5. Interestingly, there are subsets of muscle development genes that are differentially expressed in either direction throughout the time series. Both Clusters 1 and 2 show GO enrichments related to muscle development and differentiation (Fig. 1C), but on closer examination this signal appears to be driven by different gene families. The muscle development signal in Cluster 2 appears to be largely driven by the expression of genes in the MEF2 family (*Mef2c*, *Mef2d*), while the Cluster 1 genes contributing strongly to this signal are myogenic regulatory factors (*Myf5*, *Myod1*, *Myog*). Because these genes are not widely studied in the tendon and were not expected to be expressed, we sought to replicate these results using alternative techniques for assessing gene expression. RT-qPCR on *Myf5* and *Myod1* using tendon RNA collected from a new cohort of *Scx-Cre;TdTm* mice, the same strain used to generate the RNA-seq libraries (see Methods; Supp. Fig.3B, C) replicated this result showing downregulation of both genes from early to late time points. As *Scx*-lineage cells were found to fuse with myofibers and contribute to muscle (40), the detection of muscle-related transcription factors in our datasets may result from either the collection of these fibroblast-muscle cells or from direct contamination from muscle. Unlike Clusters 1 and 2, Cluster 5 is enriched for genes involved in regulating the assembly of contractile elements in striated muscle cells, specifically actin (e.g., *Acta1*, *Actn3*, *Tmod1*, *Tmod4*) and titin (e.g., *Tcap*). Although not enriched for muscle-specific processes, Cluster 4 does contain many genes involved in the regulation of the actin cytoskeleton (e.g., *Acta2*).

### ATAC-seq identifies accessible chromatin regions in tendon cells

To map chromatin accessibility and putative TF-binding events throughout postnatal tendon development, we performed ATAC-seq on 5,000 FACS-sorted *Scx*-lineage (*Scx-Cre;TdTom*^+^) cells collected from tendons at the same six time points described above. Due to insufficiencies in sequenced library quality, we had to discard both P21 replicates from the analysis. Because Tn5 transposase is known to preferentially cut certain sequences (41), we performed a control ATAC assay on naked, or genomic, DNA and sequenced this library along with those from each time point. We controlled for Tn5 sequence preference and other library preparation artifacts by filtering reads found in the control library from the data set for all downstream analyses (see Methods). All sequenced libraries are enriched for insert sizes < 250 bp, which indicates a large number of nucleosome free (<100 bp) and mono-nucleosomal (180-247 bp) regions (42). Additionally, biological replicates for each time point are well correlated (Pearson correlation > 0.8).

Localized regions of accessible chromatin (peaks) were identified for each replicate using the MACS2 peak calling algorithm (43) and a consensus peak set for all replicates was defined using the DiffBind package (44) in R yielding 67,438 consensus peaks that are accessible in both replicates of at least one time point. To determine differential accessibility of peaks over time, reads in peaks were counted for each biological replicate (see Methods) and the DESeq2 framework was implemented in DiffBind (see Methods). A PCA shows that the time points separate well along PC1, which explains 81% of the variance in the data set (Fig. 2B). However, similar to the transcriptomic data discussed above, P28 and P35 fail to fully separate along PC1 suggesting minimal differences between these two time points. This is supported by the differential accessibility analysis, which finds only 8 differentially accessible (DA) peaks between P28 and P35. In contrast to the relative similarities between the two latest time points, 8,358 peaks were found to be DA at some point between P0 and P35 (adj. p < 0.05). The majority of these DA peaks are located distal to known transcription start sites (TSS) in non-coding regions of the genome: ∼15% of DA peaks are found within the promoter of a gene and ∼34% fall in a distal intergenic region, while less than 2% of DA consensus peaks are located within a known exon (Fig. 2C, D). These results indicate that ATAC-seq identifies dynamic chromatin accessibility in non-coding regions of the mouse genome during postnatal tendon development.

**Figure 2:**
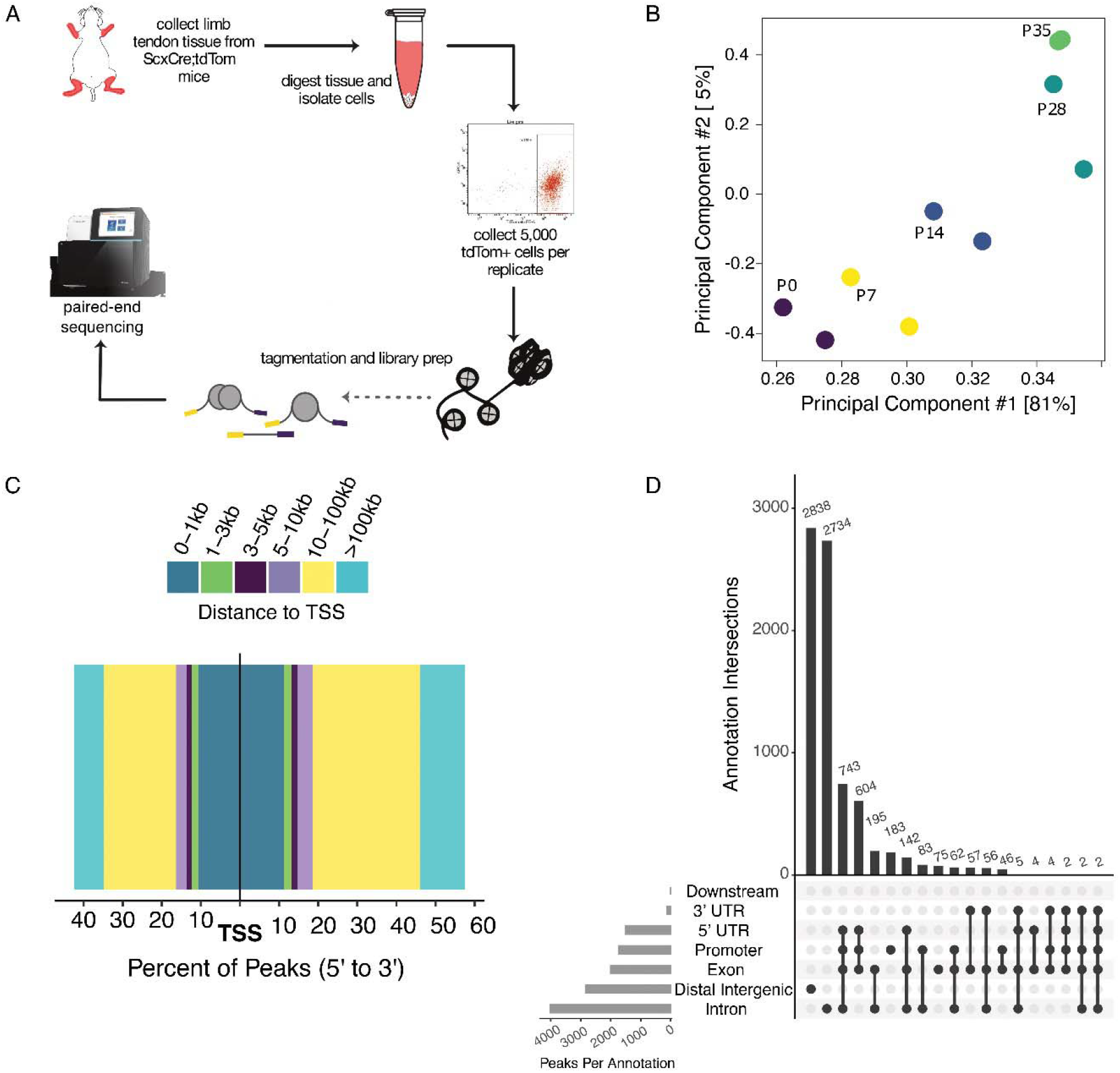
ATAC-seq of 5,000 sorted *Scx*-lineage cells detects differential chromatin accessibility during postnatal tendon development. A) Schematic of cell isolation and library preparation from mouse tendons. B) PCA of normalized counts of ATAC-seq reads in peaks shows primary separation of timepoints along PC1. Early (P0 and P7), middle (P14), and late (P28 and P35) timepoints are further separated along PC 2. C) Bar plot showing distribution of consensus ATAC-seq peaks around TSS genome-wide. Peaks are binned and colored by distance from TSS. The percent of consensus peaks in each bin is plotted along the x axis in a 5’ to 3’ orientation, i.e. bars to the left indicate peaks upstream from a TSS and those to the right are downstream. D) Upset plot of the distribution and intersection of genomic annotations assigned to consensus ATAC-seq peaks showing that the majority of peaks are found in non-coding regions.

### Integration of ATAC-seq and RNA-seq data

To test whether any of these DA regions could harbor functionally relevant *cis*-regulatory elements, we first examined the relationship between genome-wide chromatin accessibility and gene expression measured via RNA-seq across postnatal tendon development. ATAC-seq peaks were assigned to the nearest gene and Pearson correlation was computed between peak accessibility and gene expression for each peak-gene pair. Of the 6,193 DA peak-gene pairs for which gene expression was detected throughout the time series, 2,180 (∼35%) were found to exhibit a positive correlation between accessibility and expression (Pearson correlation > 0.5) indicating potential transcriptional activation (enhancer) activity by these DA peaks (Fig. 3B). Meanwhile, 1335 (∼21%) demonstrated a negative correlation (Pearson correlation < −0.5), potentially indicating that these non-coding regions may behave as repressive elements (Fig. 3B). Although we are interested in general transcriptional regulation throughout postnatal growth, the relationship between transcriptional activators and their targets is more straightforward. Because little is known about *cis*-regulatory control during tendon growth in general, we chose to focus on potential enhancer regions in downstream analyses. We applied the PAM algorithm (45) to the subset of positively correlated peak-gene pairs based on their accessibility and expression over time, which revealed three primary patterns of accessibility partitioned into five main clusters (Fig. 3C). Similar to the transcriptomic clustering results, the majority of these peaks and their assigned genes show either monotonic increasing or decreasing accessibility and expression over time. Interestingly, this co-clustering analysis also found a group of peak-gene pairs that exhibit a specific, coordinated increase at P7 (Fig. 3C), similar to the pattern reflected in Cluster 4 (expression downregulation over time) from the transcriptomic analysis (Fig. 1C).

**Figure 3:**
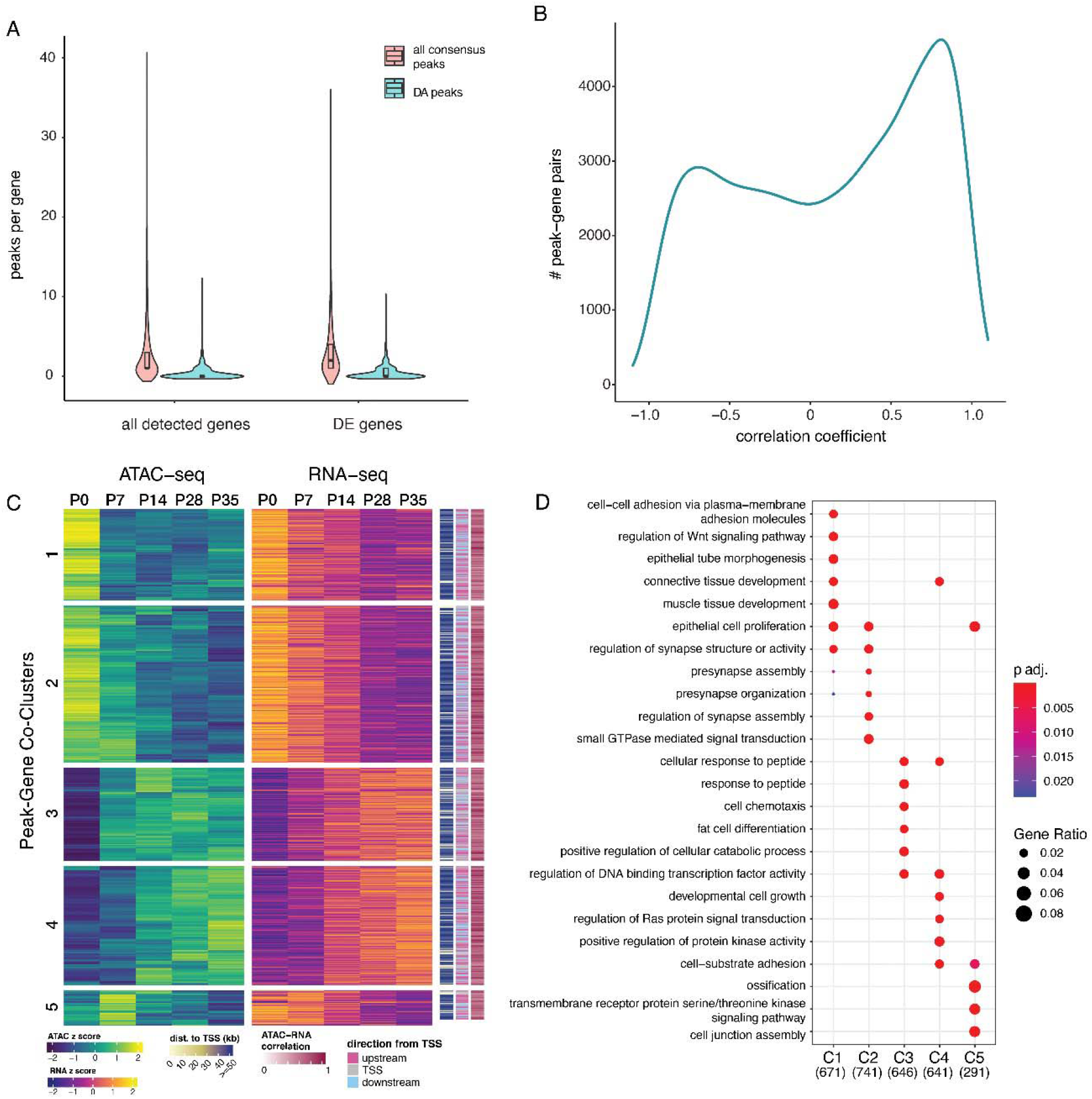
Integrative analysis of transcriptomics and chromatin accessibility reveals putative regulatory information about the postnatal transition in tendon cells. A) Violin plot showing the distribution of the number of ATAC-seq peaks assigned to a given gene. While most genes (either merely detected or DE) had ≥1 associated consensus peak, fewer genes had ≥1 associated DA peak. B) Density distribution of correlation coefficients calculated for gene expression and chromatin accessibility over time for all peak-gene pairs. C) Heatmap of ATAC-seq and RNA-seq peak-gene co-clusters determined using the PAM algorithm. Z-scores were calculated from log2(x +1)-transformed, quantile normalized CPM. Heatmap annotations are (from left to right): absolute distance of the peak from the nearest TSS; direction from TSS, i.e. upstream, downstream, or overlapping with the TSS annotation; correlation score (r2) for the peak-gene pair based on ATAC-seq and RNA-seq normalized counts across the time series. D) Dot plot depicting the top five enriched GO categories for each cluster. Bubble color represents the adjusted p value for the enrichment and bubble size represents the gene ratio for the GO term.

Next, we performed enrichment analyses on the integrated modules to determine whether the genes assigned to these putative enhancer regions are involved in similar functions identified based on gene expression alone. Much like the transcriptomic results discussed above, the two modules that characterize a coordinated decrease in chromatin accessibility and gene expression over time (Modules 1 and 2) are both enriched for peaks associated with Wnt signaling related genes (*Wnt2*, *Fzd7*, *Lgr4*, *Wisp1*, *Wls*; p adj. < 0.005), as well as mesenchymal cell differentiation, proliferation, and organ growth (e.g., *Yap1*, *Igf1*, *Dlk1*, *Jag1*). Interestingly, Module 1 is also enriched for peak-gene pairs associated with TGF-/3 signaling (e.g., *Bmpr1a*, *Bmp2*, *Jun*; Fig. 3), as is Cluster 5.

The two modules of integrated genomic data that represent upregulation over time (Modules 3 and 4) are both enriched for muscle-related GO terms, in keeping with the findings of the transcriptomic analyses, as well as MAPK signaling and cell migration (Fig. 3). The earlier of these modules to become active, Module 3, is also enriched for peaks near genes involved in fat and immune cell differentiation, hematopoiesis, and the regulation of reactive oxygen species (e.g., *Runx1*, *Foxo1*, *Foxo3*, *Cebpb*), with considerable overlap between these GO terms. Module 4 is more specifically enriched for cell adhesion, integrin binding, and heart morphogenesis-related terms. Among the genes that appear to be driving this cardiac signal include *Sav1* and *Smad6*, which are known negative regulators of Hippo signaling (through Yap) and TGF-/3 signaling, respectively. Module 5 is also enriched for peaks associated with genes involved in TGF-/3 signaling (Fig. 3).

### Motif analysis

Coordinated changes in accessibility and expression are suggestive of potential *cis*-regulatory activity at these non-coding loci, which is ultimately controlled by TF binding activity within these regions. Thus, we sought to identify which TFs may be capable of modulating the activity of these putative enhancers by searching for known DNA motifs within these peaks using the HOMER motif-finding algorithm (46). Multiple motifs were identified in each peak, with the majority of motifs being found within 100 bp of the peak summit (Supp. Fig. 4A) and slight variation in the distribution of the number of motifs found per peak across clusters (Supp. Fig. 4B). Next, we performed enrichment analyses to identify motifs associated with specific accessibility and expression patterns in each co-cluster of DA peaks (q < 0.05). We found a number of motif enrichments that are shared between modules that change in the same direction with different timing (i.e., Clusters 1 and 2; Clusters 3 and 4), but very little overlap between peaks that become more accessible (Clusters 3 and 4) and those that become less accessible from P0 to P35 (Clusters 1, 2, and 5; Supp. Fig. 4C).

Binding motifs for TFs in the bHLH, CTCF, Homeobox, MAD, RHD, and TEA families are more enriched at earlier time points, showing an accessibility pattern consistent with Cluster 1 (Fig. 4A). Specifically, motifs for Tead2, Tead4, and Smad3 show decreasing accessibility over time and, importantly, the genes encoding these TFs are also expressed at all examined time points (Fig. 4B and Supp. Fig. 4C). The enrichment patterns of “muscle” transcription factor motifs mirrors what we found in our transcriptomic analyses: Myf5, Myog, and Myod motifs are enriched in peaks that are more accessible early (Fig. 4B; Supp. Fig. 4C), while Mef2a motifs are enriched in peaks that become accessible later (Supp. Fig. 4C). As our ATAC-seq analysis was performed on isolated *Scx-Cre;TdTom*+ cells, it is possible that these data originate from the Scx-lineage fibroblasts that fuse with muscle during development (40).

**Figure 4:**
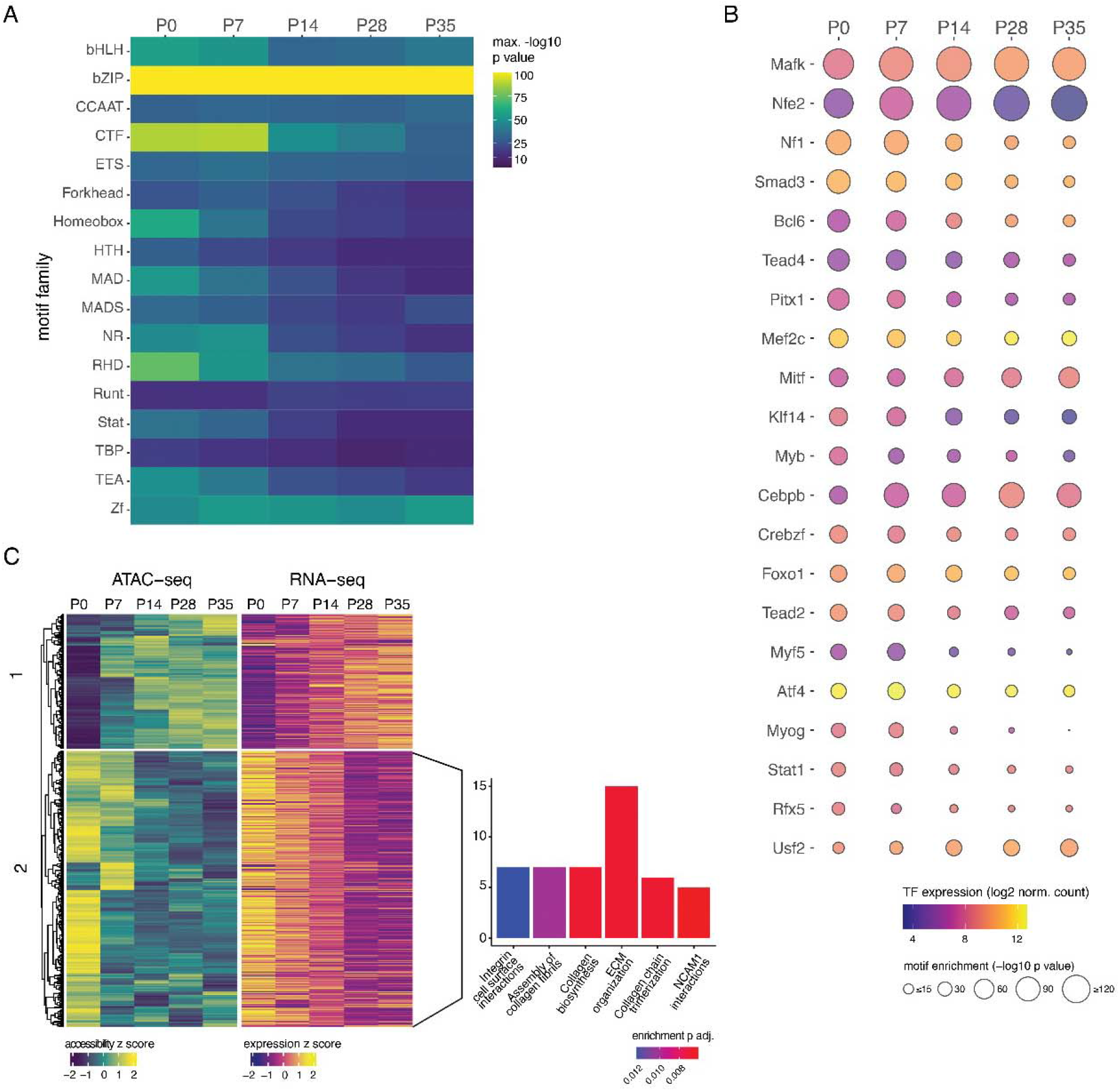
Motif analysis of differentially accessible chromatin regions identifies potential association of TEAD transcription factor with early postnatal ECM development. A) Heatmap of motif family enrichment within DA ATAC-seq peaks at each time point. B) TF motif enrichment within DA peaks and gene expression from P0 to P35. TFs were only included if they were expressed at minimum at one time point and with motif enrichment p <10-15 for at least one time point. C) Clustered heatmaps of DA ATAC-seq peaks containing TEAD motifs and expression their nearest genes. Linked bar plot shows the top 6 pathway enrichments for cluster 2; no enriched pathways were identified for cluster 1. Only positively correlated peak-gene pairs were included in this analysis.

Because the primary model of enhancer function requires interactions between the proteins bound at both the enhancer and promoter (47), it is also important to know which TFs are capable of binding the putative target promoter of a given *cis*-regulatory region. To that end, we identified TF binding motifs in the annotated proximal promoter regions of the genes assigned to these putative enhancers (positively correlated DA peak-gene pairs) and tested whether similar TF motifs, and/or those with known interactions, were enriched in both (Supp. Fig. 5A). Overall, we found fewer specific motifs enriched for each cluster (q < 0.05), but they largely reflected enrichment of the same protein families if not the same TFs (Supp. Fig.5A). Specifically, we identified enrichments of binding motifs for Tead2 (Cluster 4) and Tead4 (Clusters 1, 2, 4) in promoter-proximal regions of the putative target genes of these potential enhancers (Supp. Fig. 5A). Based on the RNA-seq data from this study, *Tead2* is significantly downregulated from P0 to P35 and an RT-qPCR analysis on independent samples verified this finding (Supp. Fig. 5B).

Because Tead2 and Tead4 binding motifs were identified as being enriched in both DA ATAC-seq peaks and their putative promoter regions in this data set, and both are expressed throughout the postnatal developmental time series, we narrowed our focus to the TEAD family of TFs. Subsetting to select only the DA peaks with TEAD family motifs and re-clustering the peak-gene pairs revealed two key patterns of accessibility and expression change from P0 to P35: upregulation (TEAD group 1) and downregulation (TEAD group 2) (Fig. 4C). We performed a pathway enrichment analysis on each set of downstream genes and found that multiple ECM and collagen related pathways were enriched in TEAD group 2 (Fig. 4C). However, no pathway enrichments were identified for TEAD group 1.

### Functional role of Yap signaling in postnatal tendon development

Given our findings that *Tead2* is downregulated in tendon tissue from P0 to P35 and Tead2 motifs are enriched in both ATAC-seq peaks that are DA at earlier timepoints and in the proximal promoters of their putative targets, we turned our attention to the broader context in which TEAD family TF binding could be functionally important during tendon postnatal development. Tead2 is a well-known co-factor of Yap/Taz; because Yap/Taz are unable to bind DNA directly, Tead co-factors play a vital role as effectors in Yap/Taz mediated Hippo signaling (48). An RT-qPCR analysis of *Yap* and *Taz* expression showed that these central signaling factors are significantly downregulated at P28 compared to P0, and both genes are differentially expressed in the RNA-seq dataset as well (Supp. Fig. 5B). Together with the results from our integrative sequencing analysis, this led us to hypothesize that Yap signaling plays an important role in postnatal tendon development and growth, specifically with respect to tendon ECM.

To test this hypothesis, we conditionally deleted *Yap* in neonatal tendon cells using *Scx-CreERt2* (Fig. 5A) and evaluated the tendon phenotype of *Yap*-conditional knockout (cKO) animals and littermate controls. We focused on Achilles tendons at P14 using multiphoton microscopy, which allowed us to assess ECM integrity via second harmonic generation (SHG) imaging. SHG signal intensity has been shown to reflect collagen structural maturity and density, thus making it a good proxy for high-collagen content ECM integrity (49). We found that the SHG intensity of *Yap*-cKO tendons was significantly lower than that of littermate controls (Fig. 5B,D; p < 0.001), indicating abnormal fibrillar collagen in the *Yap*-cKO tendons. Additionally, we found that nuclear density (number of nuclei per unit volume) was also significantly reduced in *Yap*-cKO tendons (Fig. 5B, C; p < 0.05). To assess the effects of neonatal *Yap* conditional deletion on type I collagen production, we performed RT-qPCR on *Yap*-cKO and littermate control tendons and found that *Col1a2* transcripts were significantly depleted in the *Yap*-cKO tendons (Fig. 5E; p < 0.05). This finding indicates that type I collagen production is hindered in the absence of Yap signaling, which could contribute to the observed change in SHG intensity. However, we cannot rule out the likely role of fibrillar collagen assembly and other important processes involved in the production of the tendon ECM. RT-qPCR also showed that expression of *Ki67*, a marker of cell proliferation, was significantly lower in the *Yap*-cKO samples compared to littermate controls (Fig. 5E; p < 0.05), suggesting that one possible reason for decreased nuclear density in these samples is reduced tendon cell proliferation.

**Figure 5:**
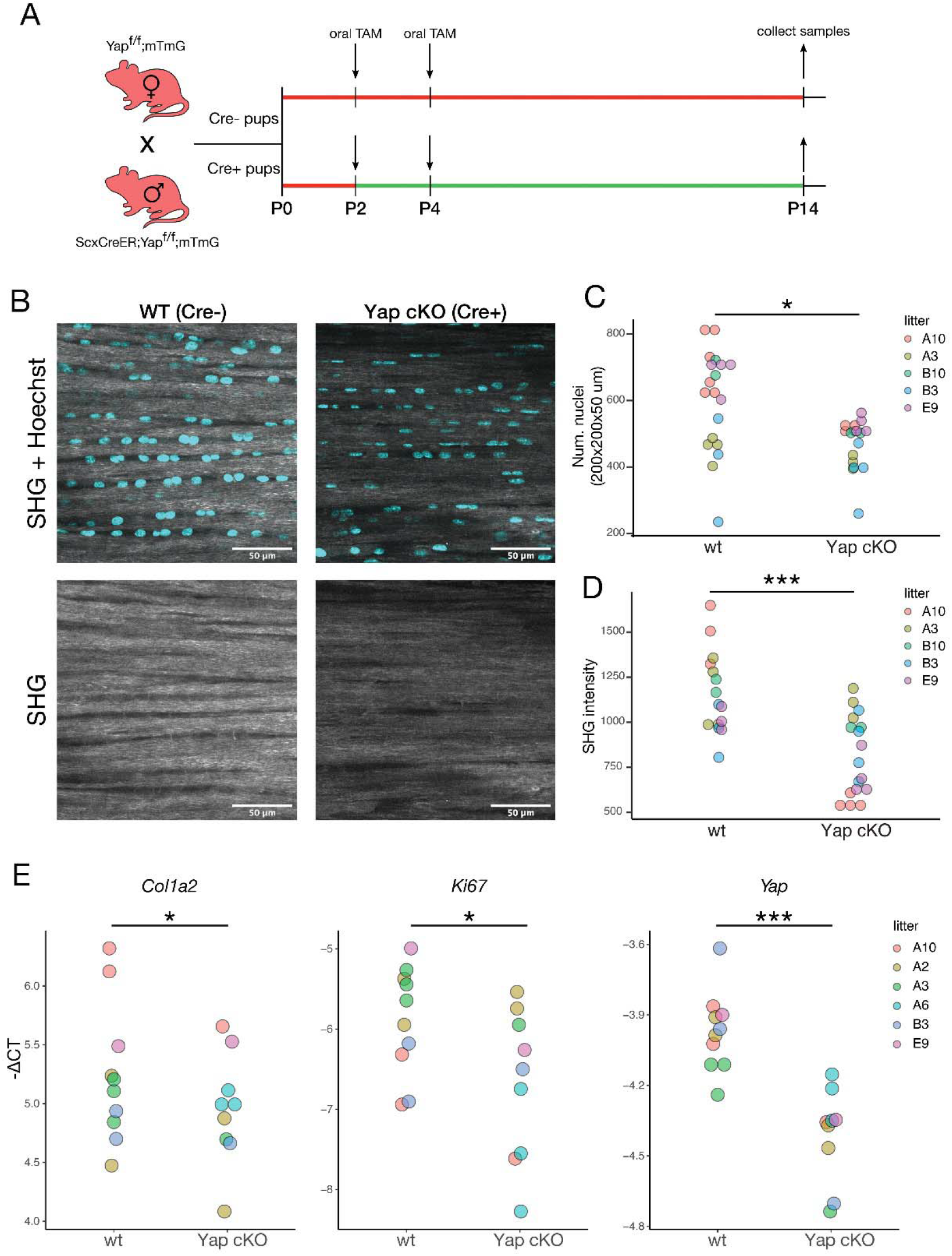
Neonatal deletion of *Yap* in *Scx*-lineage cells disrupts collagen production in the tendon during postnatal development. A) Schematic representation of mouse crosses to obtain *Yap*-cKO pups and wildtype littermates, as well as the tamoxifen dosing schedule. B) Representative multiphoton images of control (Yap^f/f^) and *Yap*-cKO (*Scx*-CreER;*Yap^f/f^*) Achilles tendon in the sagittal plane at P14. Top panel shows the merged second harmonic (SHG) and Hoechst channels; bottom shows SHG signal alone. Laser parameters were kept constant for SHG signal acquisition for control and *Yap*-cKO samples within a litter. All litter mates were imaged in the same session. Scale bars are 50 µm. C) *Yap*-cKO Achilles tendons at P14 show a decrease in nuclear density (count per unit volume) compared to controls (p < 0.05). Quantification was performed on multiphoton z-stacks. D) SHG signal intensity is significantly lower in *Yap*-cKO Achilles tendons compared to controls (p < 0.001). Intensity was measured at P14 from multiphoton z-stacks. E) RT-qPCR shows decreased expression of *Col1a2*, *Ki67*, and *Yap* in and *Yap*-cKO tendon at P14 compared to controls. −ΔCT represents expression of the target gene normalized to a reference gene (*Ppia*). * p < 0.05 0.05, ** p < 0.01, ***p < 0.001.

## Discussion

### Transcriptomics identifies differential expression of known musculoskeletal genes

Previously, little was known about the molecular changes that occur within the tendon during postnatal growth. Using a transcriptomics approach, we identified genes differentially expressed during the postnatal transition from a proliferation-based growth program to one driven by ECM expansion. Our study examines transcriptome-wide gene expression, and some previous work has examined the expression of selected target genes during this developmental period. Therefore, we can compare our results with those data to assess the validity of our findings.

### ECM genes

Most studies of postnatal tendons in mice have focused on changes to the ECM due to its important role in overall tendon tissue growth (19–21). The ECM protein collagen makes up a large portion of the dry mass of tendon (65-80%), most of which is type I collagen (38). Collagenous matrix production occurs in three phases: collagen molecule secretion and assembly into fibrils (fibrillogenesis); end-to-end fibril assembly to increase length; and lateral assembly to increase fibril diameter. We found that expression of the procollagen genes *Col1a1* and *Col1a2* peaks around P14 in accordance with our previous findings from RT-qPCR (22). Expression of procollagen genes is indicative of collagen molecule production and secretion (50, 51). It has also been previously shown that *Col3a1*, *Lum*, and *Bgn* are involved in fibril assembly and the initial stages of growth and are downregulated with age (18, 21, 52), which is supported by our data as well (Supp. Fig. 2C). In contrast to these downregulated genes, we found that *Dcn* is upregulated from P0 to P35 (p adj. < 0.01) and Fmod is upregulated from P0 to P21 (p adj. < 0.05), at which point expression appears to plateau (Supp. Fig. 2C). These genes code for proteoglycans that bind fibrillar collagens and aid in regulation of lateral fibril growth (53–55), and the expression patterns shown here largely reflect those previously reported (18, 21, 22)

### “Tendon” genes

In contrast to the dynamic changes seen among ECM genes from P0 to P35, genes known to be important for embryonic tendon development do not change significantly during postnatal development. Although they are all expressed throughout postnatal development, none of the embryonic tendon TF genes (*Scx*, *Mkx*, *Egr1*, *Egr2*) are significantly DE at any point during this time series (p adj. > 0.05). This fact does not preclude their involvement in transcriptional activation during postnatal growth, it simply shows that they are not differentially regulated themselves. We previously showed via RT-qPCR that *Scx* is significantly upregulated at P14 relative to P0 and P35, and *Mkx* is significantly downregulated by P35 compared to P0, P7, and P14 (22), contradicting our earlier findings. This discrepancy could be caused by multiple factors. First, RT-qPCR and RNA-seq measurements represent two different classes of data: RT-qPCR measures expression relative to some housekeeping gene(s) while RNA-seq measures absolute counts of transcript abundance and uses a model of transcriptome-wide expression variance to correct for biases in the data (37). Second, the RT-qPCR assay was performed on a smaller sample size (n = 3 per time point) than the RNA-seq (n = 6-8 per time point), so it is possible that variation within this small sample was not representative of the true population variance. For this reason, it is notable that the RNA-seq *Scx* measurements have far higher within-group variability compared to other tendon genes (Supp. Fig. 2A). High and low *Scx* expressing samples do not partition by sex, and none of the mice contain a *Scx* knock-out or knock-in allele in their genetic background. If such large variation in *Scx* expression is characteristic of postnatal tendon, the discrepancy between the RNA-seq results and those from the previous *Scx* RT-qPCR assay could be due to sampling bias from a highly variable population.

### “Muscle” genes

From the transcriptomics analyses, we were surprised by the number of differentially expressed genes that are typically associated with muscle. In fact, the size of these “muscle” gene cohorts in three expression modules (Clusters 1, 2, and 5) was substantial enough to influence the GO enrichment analyses (Supp. Fig. 3). Although RNA-seq Clusters 1 and 2 represent differential expression in opposite directions (downregulated vs upregulated, respectively), they are both enriched for muscle development and differentiation-related processes and functions. However, it seems that the direction of DE is partitioned by gene family. Genes in the Mef2 family of TFs (*Mef2c*, *Mef2d*) are upregulated from P0 to P35 (Cluster 2), while the myogenic regulatory factors *Myf5*, *Myod1*, and *Myog*, which are members of the bHLH TF family, are downregulated throughout postnatal development (Cluster 1). Meanwhile Cluster 5 is specifically enriched for muscle-related genes that are involved in contractile element regulation.

*Myf5* and *Myod1* play a vital role in specifying the myogenic lineage during early embryonic development (56) and in regulating muscle stem cells during adulthood (57, 58). While *Myf5* gene expression in non-muscle tissues has been reported (e.g., brown preadipocytes, (59)), *Myod1* expression is believed to be restricted to the myogenic lineage. Therefore, a possible explanation for our findings is that our samples contained cells from myotendinous regions, which include muscle cells that contained within the *Scx*-lineage (40).

Despite their association with muscle development, *Mef2* genes are expressed in a variety of other tissue types, including developing muscle (both cardiac and skeletal), embryonic chondrocytes, injured smooth muscle, and various parts of the brain (60). *Mef2c* has been shown to promote smooth muscle cell proliferation (61) and can regulate the differentiation of cardiomyocytes (62), neurons (63), and hematopoietic cells (64). Studies in Xenopus show *Mef2c* is co-expressed with *Scx* and cooperates with *Scx* to induce tendon genes (65). Both Mef2c and Mef2d have roles in regulating cytoskeletal proteins (66). Although the function of *Mef2c* and *Mef2d* transcripts in the developing tendon are unclear, it is possible that they originate from tendon cells and are involved in cytoskeletal organization of the maturing tendon cells.

### Shift in proliferative potential is correlated with transcriptome dynamics

Unsupervised clustering of significantly DE genes identified six gene co-expression modules that describe three broader patterns of expression. Cluster 1, which contains genes that are downregulated early (∼P14-P21), is highly enriched for genes involved in cell cycle. This provides transcriptomic support for our previous findings that tendon cell proliferative potential declines significantly from P0 to P21 (22). Genes involved in Wnt (Clusters 1 and 3), Hippo (Cluster 1), and Hh signaling (Cluster 3) are also overrepresented in early time points; all three of these pathways are capable of regulating cell proliferation and differentiation, and crosstalk among these signaling pathways has been previously described (reviewed in (67–69).

In addition, these pathways have been previously investigated for their role in tendon progenitor cell specification and differentiation during embryonic limb development (70). Exposure of early limb progenitor cells to Wnt maintains their proliferative abilities and increases the expression of soft connective tissue ECM genes such as *collagen 1* (*Col1a1*, *Col1a2*), *tenascin C* (*TnC*), and *Dcn* (71). Others have also shown that Wnt signaling is sufficient to inhibit chondrogenic differentiation of mesenchymal cells within the developing limb (72). More recent work has supported a role for Wnt signaling in tendon cell proliferation and suggests activation of Wnt signaling inhibits tendon gene expression in 2D cell culture (73). Sonic hedgehog (Shh) is instrumental in regulating axial tendon progenitor specification along with FGFs (74), as well as in regulating the expression of *Six1* in embryonic limb tendon (75). Meanwhile Hippo signaling and Yap have been shown to transduce mechanical signals and alter transcriptional output to influence cell fate, proliferation, and organ size control (76–79). However, this pathway has not been studied previously during tendon postnatal growth and it is currently unclear whether the rules governing its regulation in other tissues are applicable within the context of postnatal tendon. At the very least, our results are indicative of strong signatures of cell proliferation persisting into the early postnatal period that may be regulated by some combination of Wnt, Hh, and Hippo signaling.

### Changes in chromatin accessibility reflect transcriptomics and identify putative cis-regulatory regions

Overall, the results from the differential chromatin accessibility analyses reflect those from the transcriptomic analyses alone, including the myogenic signatures. As seen in the differential co-expression modules, DNA motifs for myogenic regulatory factors (e.g., Myod1) are significantly enriched within peaks that are more accessible early, while Mef2 motifs (e.g., Mef2a) are more prevalent in regions that become progressively more open during postnatal development (Supp. Fig. 4C). While this evidence from DA chromatin does not definitively indicate binding of these myogenic TFs, it is suggestive that they may play a functional role in postnatal tendon development and maturation. As our ATAC-seq was performed on a sorted population of *Scx*-lineage cells, this should have minimized the possibility that these results are contamination driven (i.e., originating from non-tendon tissues). As *Scx*-lineage cells contribute to muscle and myotendinous cells, it is likely that our analysis includes these ‘hybrid’cell types (40).

The integrative analysis demonstrates a correlation between Hippo and Wnt signaling and the proliferative period early in postnatal tendon development (Fig. 1). Peaks that are more accessible early in postnatal development tend to be associated with Wnt signaling genes (Fig. 1C) and contain AP-1 family binding motifs, including c-Jun and JunD, which are involved in the Wnt signaling pathway and cell growth (Supp. Fig. 4C, 5A) (80–82). The genes associated with these putative enhancers also contained a significant enrichment of AP-1 binding motifs in their promoter-proximal regions.

Peaks near TGF-/3 signaling factors were also found to be enriched in Modules 1 and 5, which represent early downregulation and P7-specific upregulation respectively (Supp. Fig. 6). Smad3 motifs are also significantly enriched in these peaks that become progressively less accessible over time (Modules 1, 2, and 5) (Supp. Fig. 4C). Smad3 is one of two receptor-regulated Smad proteins and is a key transducer of canonical TGF-/3 signaling (83). Increasing accessibility/expression of peak-gene pairs that negatively regulate Yap1 and TGF-/3 from P0 to P35 provides further evidence that these signaling pathways are specific to the proliferative period from P0 to P14, and that they may play a role in regulating this proliferation.

Taken together, these data suggest that some level of coordination between Wnt, TGF-/3, and Hippo signaling is involved in the transition from a highly proliferative to relatively quiescent tendon cell program. Crosstalk between the Wnt and TGF-/3 signaling pathways has been demonstrated in multiple cell types. In vascular smooth muscle cells, TGF-/3/Smad3 stimulates secretion of several Wnt ligands (Wnt2b, 4, 5a, and 9a), which promote proliferation by stabilizing /3-catenin (84). Other work in mesenchymal stem cells has shown that Tgfb1 stimulates expression of Wnt2, 4, 5a, 7a, and 10a (85, 86) during chondrogenesis, and that Smad3 and Smad4 interact with /3-catenin to activate the Wnt//3-catenin pathway in chondrogenesis (87) and osteogenic differentiation (88). It has also been shown in chondrocytes that TGF-/3 inhibits expression of *Axin1* and *Axin2* (negative regulators of Wnt) via Smad3, which inhibits TGF-/3 and promotes Wnt//3-catenin signaling (89). Additionally, previous work has shown that Smad3 can regulate tendon ECM through physical interactions with Scx and Mkx (90), both of which are stably expressed during postnatal development (Supp. Fig. 2A). Thus, it is possible that the early postnatal stage TGF-/3 signaling is indirectly regulating cell proliferation through Smad3 regulation of the matrix.

### Yap signaling in postnatal tendon cell proliferation and ECM development

Our analyses show that genes involved in the Hippo signaling pathway are differentially expressed throughout the postnatal growth period, and that progressively less accessible chromatin regions are enriched for Smad3 binding motifs (Supp. Fig. 4C). Both the RNA-seq DE analysis and RT-qPCR assays found that *Tead2*, *Yap*, and *Taz* expression decreased from P0 to P28. Furthermore, TEAD family motifs (especially Tead2 and Tead4 motifs) were found to be enriched in both DA peaks that become less accessible from P0 to P35 (Fig. 4) and their putative target gene promoters (Supp. Fig. 5A), and TEAD motif associated genes are enriched for collagen ECM related pathways. Lastly, we demonstrated that loss of Yap in neonatal tendon cells affects both the ECM and nuclear density, indicating an important role for Yap and Hippo signaling during postnatal tendon development. Although the Hippo pathway through Yap/Taz association with TEAD can regulate downstream transcriptional events, it can also interact and influence other well-known signaling pathways (91). Yap and its cofactor Taz can interact with various Wnt transcriptional activators to regulate their cytoplasmic retention in a Wnt-dependent manner. In the context of fibrosis, Yap/Taz can form complexes with Smad2/3 and Tead proteins, and the concentration of Taz influences nucleocytoplasmic shuttling of Smad proteins (reviewed in (92). Thus Yap/Taz can coordination the regulation of multiple signaling pathways.

### Limitations

Although our work provides a new resource for understanding early postnatal tendon development, the study is not without its limitations. First, RNA-seq was performed on bulk tissue, and although tendon tissue was carefully microdissected, it is possible that other cell types were present. ATAC-seq was performed on sorted *Scx*-lineage cells, however, and shows relatively good concordance with the transcriptomics results, supporting the bulk RNA-seq results. The ATAC-seq was limited to samples of 5,000 cells per biological replicate. This constrains library diversity and increases the proportion of duplicate sequenced reads, thereby limiting our power to detect more minor changes in chromatin accessibility. Ideally, to achieve this level of sensitivity, each replicate would have required at least 50,000 cells, but the intrinsic hypocellular nature of tendon tissue makes this impossible without pooling many animals. We ultimately decided that, because pooling individuals into a “biological replicate” obscures true biological variation, it would be more appropriate to perform these assays on samples that originated from a single animal. Secondly, our biological sample size for the ATAC-seq assay is limited (n = 2 per time point). Tendon tissue is difficult to dissociate without harming the cells or changing their behavior. While a larger sample size (e.g., n = 4) would have been preferable, it was simply not feasible due to the high proportion of lost samples during the dissociation and/or sorting process caused by unhealthy cells. Previous studies have performed ATAC-seq on as few as 2 biological replicates and reported reproducible results (42, 93), indicating that good quality data are possible from a limited sample number. To minimize spurious results, we constructed the consensus peak set using stringent criteria requiring a peak to be present in both biological replicates to be considered for differential accessibility; if a peak was only present in one replicate for a time point it was excluded from all downstream analyses. This likely inflates the number of false negatives in our results, but it is preferable to inflating false positives.

### Conclusions and Future Directions

Currently, we have a limited understanding of the gene regulatory networks and signaling pathways that control postnatal tendon growth and matrix maturation. As these stages also represent a turning point in regenerative to non-regenerative tendon healing and a significant transition in tendon mechanical properties, a molecular roadmap of the gene expression and genomic changes that occur during this period would be a significant advance in our ability to test new pathways in these processes. Given our findings that Wnt, Hippo, and TGF-/3 signaling appear to be downregulated in a coordinated fashion throughout postnatal development, it is possible that all three of these pathways play a role in the transition from a highly proliferative tendon cell program to a largely quiescent one. Our conditional deletion of *Yap* during the perinatal period indicates *Yap* has a role in postnatal tendon development. However, Yap deletion alone resulted in only modest, but significant, changes in tendon cell density and matrix gene expression, making it likely that its homolog, Taz, has similar functions and can compensate in its absence in regulating these processes. Targeting both Yap and Taz would further dissect the role of Hippo signaling in tendon development and maturation. Future functional studies targeting other components of this and other pathways will be vital for untangling the complicated network of interactions during tendon postnatal growth.

## Materials and Methods

### Experimental Model and Subject Details

#### Animals

*Scx-Cre* and *Scx-CreERT2* mice were provided by the Schweitzer lab (4, 94). *Gt(ROSA)26Sor^tm9(CAG-tdTomato)Hze^* (Ai9) were purchased from the Jackson Laboratory (Jax 007909) and *Gt(ROSA)26Sor^tm4(ACTB-tdTomato,-EGFP)Luo^*(referred to here as mTmG) were provided by the Rajagopal lab (REF/Jax 007676). *Yap^flox^* mice were provided by the Camargo lab (95). To obtain *Scx*-lineage cells for ATAC-seq, male *Scx-Cre^+^*mice were mated to female mice positive for the Ai9 Cre reporter allele to generate the *Scx-Cre;Rosa:TdTomato* mouse line (henceforth referred to as *Scx-Cre;TdTm*) to mark *Scx*-lineage cells.

For the *Yap* conditional knock-out (*Yap*-cKO) experiment, male *Scx-CreERT2* mice were mated to female mice homozygous for both the *Yap^flox^* allele as well as mTmG. To perform conditional knock-out of *Yap*, all pups in each litter were given 10 µl of tamoxifen in corn oil (50 mg/ml) orally at P2 and P4. Pups were then genotyped at P10 for both *Scx-CreERT2* and floxed *Yap* alleles. An equal number of male and female mice were used for all experiments. Animals in this study were euthanized via decapitation (P0) or CO2 inhalation and subsequent cervical dislocation (P7 and older). In the *Yap*-cKO mouse model, Cre recombination in tendon cells was confirmed postmortem by visual inspection of mTmG signal in transverse tendon sections. Efficiency of floxed allele excision was evaluated by PCR using primers that amplify both wildtype and mutant (excised) regions in the *Yap* locus from tail tendon gDNA (see Supp. Table 1 for primer sequences). Finally, downregulation of *Yap* in cKO mice relative to wildtype littermates was confirmed via RT-qPCR using two sets of *Yap* primers that target different regions of the gene (see Supp. Table 1 for primer sequences). Only those mice that passed all three levels of *Yap*-cKO validation were used in subsequent analyses. Tamoxifen treated *Scx-CreERT2*-littermates served as wildtype controls.

All mice used in this study were housed at the Center for Comparative Medicine at Massachusetts General Hospital (MGH) and experiments were approved by the MGH Institutional Animal Care and Use Committee (protocol #2013N000062).

### Method Details

See (Supp. Fig. 7) for an overview of the RNA isolation and library preparation methods for RNA-seq and ATAC-seq.

#### RNA Isolation

Extraction of intact total RNA from whole tendons was performed as previously described (22, 96). Briefly, distal hind limb and forelimb tendons were dissected from mice at weekly time points between P0 and P35 and submerged in cold TRIzol (Invitrogen 15596026) immediately following euthanasia. Multiple tendons from a single animal were collected in the same 1.5 mL tube containing 500 µl to 1mL of TRIzol and high impact zirconium 1.5 mm beads (Benchmark D1032-15). The volume of TRIzol used was dependent on the size of the sample. Tendons were first roughly chopped with clean microdissection scissors and then homogenized in two 180-second bouts of bead beating at 50 Hz in a BeadBug microtube homogenizer (Benchmark). Following homogenization, samples were stored at −80 °C until extraction using a combination of the TRIzol-chloroform method (97) and Zymo Direct-Zol system (Zymo Research R2050) with an on-column DNaseI digestion (Zymo Research E1010). RNA purity and quality were evaluated using spectrophotometry (NanoDrop 2000c, Thermo Scientific) and capillary electrophoresis (2100 Bioanalyzer and TapeStation 2200, Agilent), respectively. Concentration of each sample was measured via fluorometric quantitation (Qubit HS RNA assay, Invitrogen Q32852) and the final RNA product was stored at −80 °C.

#### RNA-seq library preparation and sequencing

RNA samples were excluded from RNA-seq experiments if RIN < 6.7 (threshold determined empirically), sample purity measures (260/280 and 260/230) were poor, and/or if the sample contained insufficient RNA for optimal library preparation. Between 7 and 9 biological replicates (i.e., tendon RNA from independent mice) per time point passed these quality measures, yielding a total of 46 samples spanning 6 time points. To minimize batch effects, all RNA-seq library preparation steps were completed in a single batch using the Apollo 324 NGS library prep system (IntegenX) and the PrepX mRNA library protocol (Takara Bio) in the Harvard Bauer Core Facility. First, mRNA was isolated from 1 µg total RNA by polyA selection (PrepX polyA 48, Takara Bio 640098) and checked for rRNA contamination using the mRNA 2100 Bioanalyzer chip and protocol (Agilent). The remaining mRNA for each sample was then reverse transcribed and purified using PrepX chemistry (PrepX mRNA 48, Takara Bio 640097). The resulting cDNA libraries were uniquely barcoded and amplified with 11 PCR cycles and then multiplexed into three pools of 16 samples for sequencing. Single end 75 bp reads were sequenced on an Illumina NextSeq 500 using a High-Output 75-cycle kit in the Harvard Bauer Core Facility. Sequenced libraries that achieved at least 20 million reads were used for downstream analyses.

#### Tendon cell isolation and FACS sorting for ATAC-seq

Distal hind limb and forelimb tendons were dissected from *Scx-Cre;TdTm^+^* mice at weekly time points from P0 to P35. Tendons were collected in a digestion buffer containing 0.2% collagenase type II (Worthington L5004176) in DMEM (Gibco 11965-092), coarsely chopped, and incubated in a shaking water bath at 37C. Midway through the incubation the digestion media was spiked with 0.2% collagenase type I (Gibco 17100-017) and 0.4% dispase (Gibco 17105-041). The digested tendons were then gently manually dissociated with a 20G needle and passed through a 30 µm pre-separation filter (MACS 130-041-407) to collect the cells. Tendon cells were washed with 10% horse serum in Hams F10 media (Gibco 11550-043) and finally filtered through the cell strainer cap of a FACS tube (Falcon 352235) for flow cytometry and cell sorting. Tendon cell suspensions were incubated with DAPI immediately before sorting for live/dead exclusion. TdTomato (TdTm) FACS gates were defined based on control TdTm^+^ and TdTm^−^ tendon cells that were collected and processed in parallel with the samples of interest. 5,000 TdTm^+^ cells were collected per replicate in 5% FBS/PBS. Control samples of 50 ng of naked DNA (i.e., DNA free of histones and DNA-binding proteins) from mice were also subjected to tagmentation and ATAC-seq library preparation in parallel with the tendon samples.

#### ATAC-seq library preparation and sequencing

Cells were pelleted and washed in clean 1x PBS. Transposition was performed using the Fast-ATAC protocol (98) with the following changes: 2 µl of Tn5 transposase (TDE1 from Illumina FC-121-1030) was used in the transposition reaction and the reactions were incubated at 37 °C in a shaking water bath for 35 minutes. Transposed DNA was purified using an Omega MicroElute Cleanup kit (Omega D6296) and eluted in 15 µl nuclease free water. The optimal number of PCR cycles for each library was determined via a qPCR side reaction (99). The purified transposed fragments were then PCR amplified and barcoded as described in Buenrostro et al. (99), followed by bead purification (Omega M1386-01). Library quality was examined via capillary electrophoresis (2100 Bioanalyzer and TapeStation 2200, Agilent) and libraries were quantified using the KAPA library quantification system for qPCR (KAPA KK4824). ATAC libraries were multiplexed and paired-end 42 bp reads were sequenced on an Illumina NextSeq 500 using a High-Output 75-cycle kit in the Harvard Bauer Core Facility.

#### RT-qPCR

Tendons were collected from *Scx-Cre;TdTm^+^* mice (n = 4 per time point; 20 total) and RNA was extracted as described above. Because minimal differences were found between P28 and P35 in the transcriptomics analysis, we stopped collection at P28 for RT-qPCR validation. 1 µg of total RNA from each sample was reverse transcribed using the SSIV first strand synthesis system with oligo(dT)20 primers (Thermo Fisher 18091050). SYBR green (PowerUp SYBR, Applied Biosystems A25742) qPCR assays were conducted in technical triplicate using 10 ng cDNA per reaction (final cDNA concentration = 0.8 ng/µl). Samples were amplified with target-specific primers (see Supp. Table 1 for primer sequences) using a LightCycler 480 II real time PCR system (Roche Diagnostics) as previously described (22).

#### Multiphoton microscopy

Following euthanasia, mouse hind limbs were fixed in 4% PFA overnight at 4°C. Fixed Achilles tendons were dissected from the limb, incubated in Hoechst 33258 (ThermoFisher Cat# H3569) nuclear stain (1:1000) for 2 hours at room temperature, rinsed in PBS, and mounted for whole mount multiphoton microscopy. Three Achilles samples were imaged per genotype (total n = 6) and images of at least different 3 regions of interest (ROI) within the tendon midsubstance were collected per sample. Laser parameters for each litter were calibrated on wildtype samples and used to image both control and *Yap*-cKO tendons to enable direct comparison of second harmonic intensity within a litter. Hoechst and second harmonic signal were acquired in 100 µm z stacks (0.5 µm step size) on an Olympus FluoView (FVMPE-RS) multiphoton laser scanning microscope using a 25X water immersion lens (XLPlan N 25X WMP) in the MGH Center for Regenerative Medicine Multiphoton Microscopy Core.

#### Multiphoton image analysis

All multiphoton image analysis was conducted using FIJI software (100) with custom macros. Average second harmonic intensity was computed for each slice in the z stack, from which the maximum intensity was determined. Nuclei were counted within a standardized volume (200 µm x 200 µm x 50 µm).

### Quantification and Statistical Analysis

#### RNA-seq Data Processing and Differential Expression Analysis

Sequenced RNA-seq reads were demultiplexed, followed by quality filtering and trimming using TrimGalore (101). After quality filtering, the final sample size was between 6 and 9 biological replicates per time point. A Salmon transcript index was built from the Ensembl mouse transcriptome (GRCm38v91) and transcripts were then quantified from quality-trimmed reads using Salmon (102) in mapping-based mode with the `--seqBias` and `-- gcBias` flags enabled. When used, these options allow the Salmon algorithm to learn and correct for sequence specific and GC biases, respectively, in the data. Gene-level counts were then calculated with TxImport (103). Genes with consistently low counts were excluded from the data set using automatic independent filtering implemented in DESeq2 (37, 104). Differential expression analyses were conducted using DESeq2 (37). We defined two negative binomial generalized linear models of gene expression using RIN as a blocking factor:

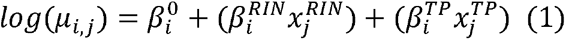

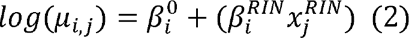

where *i* = *gene*, *j* = *sample*. The full model (Equation (1)) includes both RIN and time point as predictor variables whereas RIN is the only predictor in the reduced model (Equation (2)). Using a likelihood ratio test (LRT) we compared the two models to identify significant genes that are explained by time point, but not RIN, in order to computationally account for any effects of RNA integrity on the results. The significance threshold for differential expression was set at p adj. < 0.05.

#### ATAC-seq Data Processing and Differential Accessibility Analysis

Demultiplexed paired end reads were trimmed using NGmerge (105) and aligned to the mouse genome using Bowtie2 (106). Read pairs mapping to the mitochondrial genome were removed using removeChrom (https://github.com/jsh58/harvard) and PCR duplicates were filtered with the `dedup` tool in bamUtil (107). Peaks were called with MACS2 in paired end BAM mode with the options `-f BAMPE --nolambda --keep-dup all`. Any peaks falling in the ENCODE defined blacklist for mm10 (108) and peaks called from the naked DNA control were deemed spurious and removed using bedtools (109). Two successful biological replicates were obtained for all time points except P21, which was excluded from all analyses.

We defined a consensus peak set using the DiffBind R package (44) based on stringent requirements. In order for a peak to be included in the consensus set it had to be present in both biological replicates for at least one time point. Peaks that did not meet this criterion were deemed irreproducible and were excluded from all downstream analyses. We also constrained consensus peak widths to 500 bp. ATAC-seq peaks were annotated to genomic features in R using the ChIPseeker package (110) and were assigned to the nearest gene within a 100 kb window, defined as ± 50 kb from the consensus peak summit. DiffBind was used to count reads in these consensus peaks and compute peak differential accessibility (DA) using the underlying DESeq2 framework. The significance threshold was set at p adj. < 0.05.

#### Clustering and Functional Enrichment

Prior to all clustering analyses, counts of significantly DE genes and DA peaks were normalized using the DESeq2 framework (median of ratios method) and scaled to library size to produce log2 counts per million (CPM). These counts were quantile-normalized and z-transformed. Pearson distance matrices were then calculated for the entire gene or peak set. Clusters were computed in R (R Core Team, 2019) based on this distance matrix and the normalized/transformed counts using the Partitioning Around Medoids (PAM) algorithm (45) implemented in the ClusterR package (111) with the `fuzzy` option enabled. Heatmaps of clusters were constructed using the ComplexHeatmap package (112) and genes (rows) within each PAM module were subclustered via average linkage hierarchical clustering to aid visualization. Functional enrichment analyses of peak and gene clusters were conducted in R using the clusterProfiler package (113) with significance thresholds set at p adj. < 0.01 and q < 0.05. The Benjamini Hochberg method was used for p value adjustment.

#### Integrative Analyses

In order to compare the patterns of gene expression and chromatin accessibility, DA peaks were clustered as described above (Methods, Clustering and Functional Enrichment) into five modules. Next, a matrix of normalized counts for each peak’s assigned gene was generated and this matrix was aligned with the matrix of normalized DA peak counts. We computed the Pearson correlation coefficient for each peak-gene pair using the ‘lineup’ R package (114). For co-clustering of expression and accessibility patterns, the data set was filtered for a Pearson correlation coefficient > 0.5. This matrix of positively correlated of peak-gene pairs were re-clustered into five modules with the PAM algorithm. Gene lists for the final integrated clusters were used for functional analyses as described above (Methods, Clustering and Functional Enrichment). For each integrative cluster, known motifs were identified within peaks and putative target gene promoters using the HOMER motif finding algorithm (46). Per cluster motif enrichment calculations based these on these motif predictions were facilitated by the ‘marge’ (115) and ‘valr’ (116) packages for R.

#### RT-qPCR statistics

Cycle threshold (*C^T^*) values for all targets were normalized to *Gapdh* and the delta delta Ct method (117) was used to calculate relative gene expression for visualization. Relative expression is visualized as 2*^ΔΔCt^* ± standard deviation. Statistics were computed on the delta Ct values. Statistical differences among the time points were investigated using a Kruskal-Wallis rank sum test followed by a Dunn test with Benjamini-Hochberg correction to test for specific differences among pairs of time points. Statistics were performed on the delta Ct values (normalized to *Gapdh*) for all target genes and samples (n = 4 biological replicates per time point). R statistical software (R Core Team, 2019) was used for all RT-qPCR data analysis, statistics, and visualization. Statistical analysis was performed using the implementation of the Kruskal-Wallace test in ‘stats’ (version 3.5.3) (R Core Team, 2019) and the Dunn test from ‘FSA’ (version 0.8.25; (118)).

#### Multiphoton image analysis statistics

For both SHG intensity and nuclear density, differences between the controls and Yap-cKOs were assessed with linear mixed-effects models (LMMs) using the lmer function from lme4. The LMMs included both litter and sample as random effects.

#### Software and computing environment

Processing and large-scale analysis of sequencing data was performed on the Harvard Odyssey computing cluster (centOS7). Python programs were run in Python version 2.7.12. R programs were run in R version 3.5.3 (R Core Team, 2019) and RStudio version 1.0.143 (RStudio Team, 2015). Data analysis and visualization in R was assisted by R packages included in the Tidyverse collection (119), ‘ggplot2’ (120), and ‘viridis’ (121).

#### Data availability

Raw and processed RNA-seq and ATAC-seq data files have been deposited in NCBI’s Gene Expression Omnibus (GEO) and are available through GEO Series accession number GSE272888.

## References

1. Montgomery RD. Healing of muscle, ligaments, and tendons. Semin Vet Med Surg Small Anim. 1989;4(4):304–11.

2. Beredjiklian PK, Favata M, Cartmell JS, Flanagan CL, Crombleholme TM, Soslowsky LJ. Regenerative versus reparative healing in tendon: a study of biomechanical and histological properties in fetal sheep. Ann Biomed Eng. 2003;31(10):1143–52.

3. Gomoll AH, Katz JN, Warner JJ, Millett PJ. Rotator cuff disorders: recognition and management among patients with shoulder pain. Arthritis Rheum. 2004;50(12):3751–61.

4. Schweitzer R, Chyung JH, Murtaugh LC, Brent AE, Rosen V, Olson EN, et al. Analysis of the tendon cell fate using Scleraxis, a specific marker for tendons and ligaments. Development. 2001;128(19):3855–66.

5. Lejard V, Blais F, Guerquin MJ, Bonnet A, Bonnin MA, Havis E, et al. EGR1 and EGR2 involvement in vertebrate tendon differentiation. J Biol Chem. 2011;286(7):5855–67.

6. Liu Y, Schwartz AG, Birman V, Thomopoulos S, Genin GM. Stress amplification during development of the tendon-to-bone attachment. Biomech Model Mechanobiol. 2014;13(5):973–83.

7. Brandau O, Meindl A, Fassler R, Aszodi A. A novel gene, tendin, is strongly expressed in tendons and ligaments and shows high homology with chondromodulin-I. Dev Dyn. 2001;221(1):72–80.

8. Shukunami C, Oshima Y, Hiraki Y. Molecular cloning of tenomodulin, a novel chondromodulin-I related gene. Biochem Biophys Res Commun. 2001;280(5):1323–7.

9. Docheva D, Hunziker EB, Fassler R, Brandau O. Tenomodulin is necessary for tenocyte proliferation and tendon maturation. Mol Cell Biol. 2005;25(2):699–705.

10. Subramanian A, Schilling TF. Tendon development and musculoskeletal assembly: emerging roles for the extracellular matrix. Development. 2015;142(24):4191–204.

11. Ignotz RA, Massague J. Transforming growth factor-beta stimulates the expression of fibronectin and collagen and their incorporation into the extracellular matrix. J Biol Chem. 1986;261(9):4337–45.

12. Montesano R, Orci L. Transforming growth factor beta stimulates collagen-matrix contraction by fibroblasts: implications for wound healing. Proc Natl Acad Sci U S A. 1988;85(13):4894–7.

13. Pryce BA, Watson SS, Murchison ND, Staverosky JA, Dunker N, Schweitzer R. Recruitment and maintenance of tendon progenitors by TGFbeta signaling are essential for tendon formation. Development. 2009;136(8):1351–61.

14. Havis E, Bonnin MA, Esteves de Lima J, Charvet B, Milet C, Duprez D. TGFbeta and FGF promote tendon progenitor fate and act downstream of muscle contraction to regulate tendon differentiation during chick limb development. Development. 2016;143(20):3839–51.

15. Maeda T, Sakabe T, Sunaga A, Sakai K, Rivera AL, Keene DR, et al. Conversion of mechanical force into TGF-beta-mediated biochemical signals. Curr Biol. 2011;21(11):933–41.

16. Mendias CL, Gumucio JP, Lynch EB. Mechanical loading and TGF-beta change the expression of multiple miRNAs in tendon fibroblasts. J Appl Physiol (1985). 2012;113(1):56–62.

17. Tan GK, Pryce BA, Stabio A, Brigande JV, Wang C, Xia Z, et al. Tgfbeta signaling is critical for maintenance of the tendon cell fate. Elife. 2020;9.

18. Ansorge HL, Adams S, Birk DE, Soslowsky LJ. Mechanical, compositional, and structural properties of the post-natal mouse Achilles tendon. Ann Biomed Eng. 2011;39(7):1904–13.

19. Ahmadzadeh H, Connizzo BK, Freedman BR, Soslowsky LJ, Shenoy VB. Determining the contribution of glycosaminoglycans to tendon mechanical properties with a modified shear-lag model. J Biomech. 2013;46(14):2497–503.

20. Kalson NS, Lu Y, Taylor SH, Starborg T, Holmes DF, Kadler KE. A structure-based extracellular matrix expansion mechanism of fibrous tissue growth. Elife. 2015;4.

21. Ezura Y, Chakravarti S, Oldberg A, Chervoneva I, Birk DE. Differential expression of lumican and fibromodulin regulate collagen fibrillogenesis in developing mouse tendons. J Cell Biol. 2000;151(4):779–88.

22. Grinstein M, Dingwall HL, O’Connor LD, Zou K, Capellini TD, Galloway JL. A distinct transition from cell growth to physiological homeostasis in the tendon. Elife. 2019;8.

23. Ansorge HL, Hsu JE, Edelstein L, Adams S, Birk DE, Soslowsky LJ. Recapitulation of the Achilles tendon mechanical properties during neonatal development: a study of differential healing during two stages of development in a mouse model. J Orthop Res. 2012;30(3):448–56.

24. Howell K, Chien C, Bell R, Laudier D, Tufa SF, Keene DR, et al. Novel Model of Tendon Regeneration Reveals Distinct Cell Mechanisms Underlying Regenerative and Fibrotic Tendon Healing. Sci Rep. 2017;7:45238.

25. Favata M, Beredjiklian PK, Zgonis MH, Beason DP, Crombleholme TM, Jawad AF, et al. Regenerative properties of fetal sheep tendon are not adversely affected by transplantation into an adult environment. J Orthop Res. 2006;24(11):2124–32.

26. Porrello ER, Mahmoud AI, Simpson E, Hill JA, Richardson JA, Olson EN, et al. Transient regenerative potential of the neonatal mouse heart. Science. 2011;331(6020):1078–80.

27. Xin M, Kim Y, Sutherland LB, Murakami M, Qi X, McAnally J, et al. Hippo pathway effector Yap promotes cardiac regeneration. Proc Natl Acad Sci U S A. 2013;110(34):13839–44.

28. Screen HR, Berk DE, Kadler KE, Ramirez F, Young MF. Tendon functional extracellular matrix. J Orthop Res. 2015;33(6):793–9.

29. Havis E, Bonnin MA, Olivera-Martinez I, Nazaret N, Ruggiu M, Weibel J, et al. Transcriptomic analysis of mouse limb tendon cells during development. Development. 2014;141(19):3683–96.

30. Liu H, Xu J, Liu CF, Lan Y, Wylie C, Jiang R. Whole transcriptome expression profiling of mouse limb tendon development by using RNA-seq. J Orthop Res. 2015;33(6):840–8.

31. Kohler J, Popov C, Klotz B, Alberton P, Prall WC, Haasters F, et al. Uncovering the cellular and molecular changes in tendon stem/progenitor cells attributed to tendon aging and degeneration. Aging Cell. 2013;12(6):988–99.

32. Jelinsky SA, Archambault J, Li L, Seeherman H. Tendon-selective genes identified from rat and human musculoskeletal tissues. J Orthop Res. 2010;28(3):289–97.

33. Peffers MJ, Fang Y, Cheung K, Wei TK, Clegg PD, Birch HL. Transcriptome analysis of ageing in uninjured human Achilles tendon. Arthritis Res Ther. 2015;17:33.

34. Harvey T, Flamenco S, Fan CM. A Tppp3(+)Pdgfra(+) tendon stem cell population contributes to regeneration and reveals a shared role for PDGF signalling in regeneration and fibrosis. Nat Cell Biol. 2019;21(12):1490–503.

35. De Micheli AJ, Swanson JB, Disser NP, Martinez LM, Walker NR, Oliver DJ, et al. Single-cell Transcriptomic Analyses Identifies Extensive Heterogeneity in the Cellular Composition of Mouse Achilles Tendons. Am J Physiol Cell Physiol. 2020.

36. Grinstein M, Tsai SL, Montoro D, Dingwall HL, Zou K, Sade-Feldman M, et al. A quiescent resident progenitor pool is the central organizer of tendon healing. bioRxiv. 2022:2022.02.02.478533.

37. Love MI, Huber W, Anders S. Moderated estimation of fold change and dispersion for RNA-seq data with DESeq2. Genome Biol. 2014;15(12):550.

38. Kannus P. Structure of the tendon connective tissue. Scand J Med Sci Sports. 2000;10(6):312–20.

39. Kaji DA, Howell KL, Balic Z, Hubmacher D, Huang AH. Tgfbeta signaling is required for tenocyte recruitment and functional neonatal tendon regeneration. Elife. 2020;9.

40. Esteves de Lima J, Blavet C, Bonnin MA, Hirsinger E, Comai G, Yvernogeau L, et al. Unexpected contribution of fibroblasts to muscle lineage as a mechanism for limb muscle patterning. Nat Commun. 2021;12(1):3851.

41. Green B, Bouchier C, Fairhead C, Craig NL, Cormack BP. Insertion site preference of Mu, Tn5, and Tn7 transposons. Mob DNA. 2012;3(1):3.

42. Buenrostro JD, Giresi PG, Zaba LC, Chang HY, Greenleaf WJ. Transposition of native chromatin for fast and sensitive epigenomic profiling of open chromatin, DNA-binding proteins and nucleosome position. Nat Methods. 2013;10(12):1213–8.

43. Zhang Y, Liu T, Meyer CA, Eeckhoute J, Johnson DS, Bernstein BE, et al. Model-based analysis of ChIP-Seq (MACS). Genome Biol. 2008;9(9):R137.

44. Ross-Innes CS, Stark R, Teschendorff AE, Holmes KA, Ali HR, Dunning MJ, et al. Differential oestrogen receptor binding is associated with clinical outcome in breast cancer. Nature. 2012;481(7381):389–93.

45. Kaufman LaR, P.J. Partitioning around Medoids (Program PAM). Kaufman LaR, P.J., editor. Hoboken: John Wiley & Sons, Inc.; 1990. 68–125 p.

46. Li Y, Ni P, Zhang S, Li G, Su Z. ProSampler: an ultrafast and accurate motif finder in large ChIP-seq datasets for combinatory motif discovery. Bioinformatics. 2019;35(22):4632–9.

47. Zabidi MA, Stark A. Regulatory Enhancer-Core-Promoter Communication via Transcription Factors and Cofactors. Trends Genet. 2016;32(12):801–14.

48. Battilana G, Zanconato F, Piccolo S. Mechanisms of YAP/TAZ transcriptional control. Cell Stress. 2021;5(11):167–72.

49. Williams RM, Zipfel WR, Webb WW. Interpreting second-harmonic generation images of collagen I fibrils. Biophys J. 2005;88(2):1377–86.

50. Birk DE, Nurminskaya MV, Zycband EI. Collagen fibrillogenesis in situ: fibril segments undergo post-depositional modifications resulting in linear and lateral growth during matrix development. Dev Dyn. 1995;202(3):229–43.

51. Zhang G, Young BB, Ezura Y, Favata M, Soslowsky LJ, Chakravarti S, et al. Development of tendon structure and function: regulation of collagen fibrillogenesis. J Musculoskelet Neuronal Interact. 2005;5(1):5–21.

52. Birk DE, Mayne R. Localization of collagen types I, III and V during tendon development. Changes in collagen types I and III are correlated with changes in fibril diameter. Eur J Cell Biol. 1997;72(4):352–61.

53. Hedbom E, Heinegard D. Binding of fibromodulin and decorin to separate sites on fibrillar collagens. J Biol Chem. 1993;268(36):27307–12.

54. Rada JA, Cornuet PK, Hassell JR. Regulation of corneal collagen fibrillogenesis in vitro by corneal proteoglycan (lumican and decorin) core proteins. Exp Eye Res. 1993;56(6):635–48.

55. Vogel KG, Paulsson M, Heinegard D. Specific inhibition of type I and type II collagen fibrillogenesis by the small proteoglycan of tendon. Biochem J. 1984;223(3):587–97.

56. Rudnicki MA, Schnegelsberg PN, Stead RH, Braun T, Arnold HH, Jaenisch R. MyoD or Myf-5 is required for the formation of skeletal muscle. Cell. 1993;75(7):1351–9.

57. Ustanina S, Carvajal J, Rigby P, Braun T. The myogenic factor Myf5 supports efficient skeletal muscle regeneration by enabling transient myoblast amplification. Stem Cells. 2007;25(8):2006–16.

58. Asakura A, Hirai H, Kablar B, Morita S, Ishibashi J, Piras BA, et al. Increased survival of muscle stem cells lacking the MyoD gene after transplantation into regenerating skeletal muscle. Proc Natl Acad Sci U S A. 2007;104(42):16552–7.

59. Timmons JA, Wennmalm K, Larsson O, Walden TB, Lassmann T, Petrovic N, et al. Myogenic gene expression signature establishes that brown and white adipocytes originate from distinct cell lineages. Proc Natl Acad Sci U S A. 2007;104(11):4401–6.

60. Pon JR, Marra MA. MEF2 transcription factors: developmental regulators and emerging cancer genes. Oncotarget. 2016;7(3):2297–312.

61. Lin Q, Lu J, Yanagisawa H, Webb R, Lyons GE, Richardson JA, et al. Requirement of the MADS-box transcription factor MEF2C for vascular development. Development. 1998;125(22):4565–74.

62. Voronova A, Al Madhoun A, Fischer A, Shelton M, Karamboulas C, Skerjanc IS. Gli2 and MEF2C activate each other’s expression and function synergistically during cardiomyogenesis in vitro. Nucleic Acids Res. 2012;40(8):3329–47.

63. Li H, Radford JC, Ragusa MJ, Shea KL, McKercher SR, Zaremba JD, et al. Transcription factor MEF2C influences neural stem/progenitor cell differentiation and maturation in vivo. Proc Natl Acad Sci U S A. 2008;105(27):9397–402.

64. Cante-Barrett K, Pieters R, Meijerink JP. Myocyte enhancer factor 2C in hematopoiesis and leukemia. Oncogene. 2014;33(4):403–10.

65. della Gaspera B, Armand AS, Sequeira I, Lecolle S, Gallien CL, Charbonnier F, et al. The Xenopus MEF2 gene family: evidence of a role for XMEF2C in larval tendon development. Dev Biol. 2009;328(2):392–402.

66. Potthoff MJ, Olson EN. MEF2: a central regulator of diverse developmental programs. Development. 2007;134(23):4131–40.

67. Kim M, Jho EH. Cross-talk between Wnt/beta-catenin and Hippo signaling pathways: a brief review. BMB Rep. 2014;47(10):540–5.

68. McNeill H, Woodgett JR. When pathways collide: collaboration and connivance among signalling proteins in development. Nat Rev Mol Cell Biol. 2010;11(6):404–13.

69. Zhao B, Li L, Guan KL. Hippo signaling at a glance. J Cell Sci. 2010;123(Pt 23):4001–6.

70. Zhu X, Zhu H, Zhang L, Huang S, Cao J, Ma G, et al. Wls-mediated Wnts differentially regulate distal limb patterning and tissue morphogenesis. Dev Biol. 2012;365(2):328–38.

71. ten Berge D, Brugmann SA, Helms JA, Nusse R. Wnt and FGF signals interact to coordinate growth with cell fate specification during limb development. Development. 2008;135(19):3247–57.

72. Hartmann C, Tabin CJ. Wnt-14 plays a pivotal role in inducing synovial joint formation in the developing appendicular skeleton. Cell. 2001;104(3):341–51.

73. Walia B, Li TM, Crosio G, Montero AM, Huang AH. Axin2-lineage cells contribute to neonatal tendon regeneration. Connect Tissue Res. 2022;63(5):530–43.

74. Brent AE, Schweitzer R, Tabin CJ. A somitic compartment of tendon progenitors. Cell. 2003;113(2):235–48.

75. Bonnin MA, Laclef C, Blaise R, Eloy-Trinquet S, Relaix F, Maire P, et al. Six1 is not involved in limb tendon development, but is expressed in limb connective tissue under Shh regulation. Mech Dev. 2005;122(4):573–85.

76. Driscoll TP, Cosgrove BD, Heo SJ, Shurden ZE, Mauck RL. Cytoskeletal to Nuclear Strain Transfer Regulates YAP Signaling in Mesenchymal Stem Cells. Biophys J. 2015;108(12):2783–93.

77. Egerbacher M, Arnoczky SP, Caballero O, Lavagnino M, Gardner KL. Loss of homeostatic tension induces apoptosis in tendon cells: an in vitro study. Clin Orthop Relat Res. 2008;466(7):1562–8.

78. Low BC, Pan CQ, Shivashankar GV, Bershadsky A, Sudol M, Sheetz M. YAP/TAZ as mechanosensors and mechanotransducers in regulating organ size and tumor growth. FEBS Lett. 2014;588(16):2663–70.

79. Schiele NR, Marturano JE, Kuo CK. Mechanical factors in embryonic tendon development: potential cues for stem cell tenogenesis. Curr Opin Biotechnol. 2013;24(5):834–40.

80. Castellazzi M, Spyrou G, La Vista N, Dangy JP, Piu F, Yaniv M, et al. Overexpression of c-jun, junB, or junD affects cell growth differently. Proc Natl Acad Sci U S A. 1991;88(20):8890–4.

81. Kan A, Tabin CJ. c-Jun is required for the specification of joint cell fates. Genes Dev. 2013;27(5):514–24.

82. Mann B, Gelos M, Siedow A, Hanski ML, Gratchev A, Ilyas M, et al. Target genes of beta-catenin-T cell-factor/lymphoid-enhancer-factor signaling in human colorectal carcinomas. Proc Natl Acad Sci U S A. 1999;96(4):1603–8.

83. Massague J, Xi Q. TGF-beta control of stem cell differentiation genes. FEBS Lett. 2012;586(14):1953–8.

84. DiRenzo DM, Chaudhary MA, Shi X, Franco SR, Zent J, Wang K, et al. A crosstalk between TGF-beta/Smad3 and Wnt/beta-catenin pathways promotes vascular smooth muscle cell proliferation. Cell Signal. 2016;28(5):498–505.

85. Tuli R, Tuli S, Nandi S, Huang X, Manner PA, Hozack WJ, et al. Transforming growth factor-beta-mediated chondrogenesis of human mesenchymal progenitor cells involves N-cadherin and mitogen-activated protein kinase and Wnt signaling cross-talk. J Biol Chem. 2003;278(42):41227–36.

86. Zhou S. TGF-beta regulates beta-catenin signaling and osteoblast differentiation in human mesenchymal stem cells. J Cell Biochem. 2011;112(6):1651–60.

87. Zhang L, Su P, Xu C, Yang J, Yu W, Huang D. Chondrogenic differentiation of human mesenchymal stem cells: a comparison between micromass and pellet culture systems. Biotechnol Lett. 2010;32(9):1339–46.

88. Yang Z, Jiang H, Chachainasakul T, Gong S, Yang XW, Heintz N, et al. Modified bacterial artificial chromosomes for zebrafish transgenesis. Methods. 2006;39(3):183–8.

89. Dao DY, Yang X, Chen D, Zuscik M, O’Keefe RJ. Axin1 and Axin2 are regulated by TGF- and mediate cross-talk between TGF- and Wnt signaling pathways. Ann N Y Acad Sci. 2007;1116:82–99.

90. Berthet E, Chen C, Butcher K, Schneider RA, Alliston T, Amirtharajah M. Smad3 binds Scleraxis and Mohawk and regulates tendon matrix organization. J Orthop Res. 2013;31(9):1475–83.

91. Totaro A, Panciera T, Piccolo S. YAP/TAZ upstream signals and downstream responses. Nat Cell Biol. 2018;20(8):888–99.

92. Piersma B, Bank RA, Boersema M. Signaling in Fibrosis: TGF-beta, WNT, and YAP/TAZ Converge. Front Med (Lausanne). 2015;2:59.

93. Gehrke AR, Neverett E, Luo YJ, Brandt A, Ricci L, Hulett RE, et al. Acoel genome reveals the regulatory landscape of whole-body regeneration. Science. 2019;363(6432).

94. Blitz E, Viukov S, Sharir A, Shwartz Y, Galloway JL, Pryce BA, et al. Bone ridge patterning during musculoskeletal assembly is mediated through SCX regulation of Bmp4 at the tendon-skeleton junction. Dev Cell. 2009;17(6):861–73.

95. Camargo FD, Gokhale S, Johnnidis JB, Fu D, Bell GW, Jaenisch R, et al. YAP1 increases organ size and expands undifferentiated progenitor cells. Curr Biol. 2007;17(23):2054–60.

96. Grinstein M, Dingwall HL, Shah RR, Capellini TD, Galloway JL. A robust method for RNA extraction and purification from a single adult mouse tendon. PeerJ. 2018;6:e4664.

97. Rio DC, Ares M, Jr., Hannon GJ, Nilsen TW. Purification of RNA using TRIzol (TRI reagent). Cold Spring Harb Protoc. 2010;2010(6):pdb prot5439.

98. Corces MR, Trevino AE, Hamilton EG, Greenside PG, Sinnott-Armstrong NA, Vesuna S, et al. An improved ATAC-seq protocol reduces background and enables interrogation of frozen tissues. Nat Methods. 2017;14(10):959–62.

99. Buenrostro JD, Wu B, Chang HY, Greenleaf WJ. ATAC-seq: A Method for Assaying Chromatin Accessibility Genome-Wide. Curr Protoc Mol Biol. 2015;109:21 9 1-9.

100. Schindelin J, Arganda-Carreras I, Frise E, Kaynig V, Longair M, Pietzsch T, et al. Fiji: an open-source platform for biological-image analysis. Nat Methods. 2012;9(7):676–82.

101. Andrews SK, F; Segonds-Pichon, A; Biggins, L; Krueger, C; Wingett, S. FastQC. 2012.

102. Patro R, Duggal G, Love MI, Irizarry RA, Kingsford C. Salmon provides fast and bias-aware quantification of transcript expression. Nat Methods. 2017;14(4):417–9.

103. Soneson C, Love MI, Robinson MD. Differential analyses for RNA-seq: transcript-level estimates improve gene-level inferences. F1000Res. 2015;4:1521.

104. Bourgon R, Gentleman R, Huber W. Independent filtering increases detection power for high-throughput experiments. Proc Natl Acad Sci U S A. 2010;107(21):9546–51.

105. Gaspar JM. NGmerge: merging paired-end reads via novel empirically-derived models of sequencing errors. BMC Bioinformatics. 2018;19(1):536.

106. Langmead B, Salzberg SL. Fast gapped-read alignment with Bowtie 2. Nat Methods. 2012;9(4):357–9.

107. Jun G, Wing MK, Abecasis GR, Kang HM. An efficient and scalable analysis framework for variant extraction and refinement from population-scale DNA sequence data. Genome Res. 2015;25(6):918–25.

108. Agarwal S, Loder SJ, Cholok D, Peterson J, Li J, Breuler C, et al. Scleraxis-Lineage Cells Contribute to Ectopic Bone Formation in Muscle and Tendon. Stem Cells. 2017;35(3):705–10.

109. Quinlan AR, Hall IM. BEDTools: a flexible suite of utilities for comparing genomic features. Bioinformatics. 2010;26(6):841–2.

110. Yu G, Wang LG, He QY. ChIPseeker: an R/Bioconductor package for ChIP peak annotation, comparison and visualization. Bioinformatics. 2015;31(14):2382–3.

111. Mouselimis L. ClusterR: Gaussian mixture models, k-means, mini-batch-kmeans, k-medoids and affinity propagation clustering. 2019.

112. Gu Z, Eils R, Schlesner M. Complex heatmaps reveal patterns and correlations in multidimensional genomic data. Bioinformatics. 2016;32(18):2847–9.

113. Yu G, Wang LG, Han Y, He QY. clusterProfiler: an R package for comparing biological themes among gene clusters. OMICS. 2012;16(5):284–7.

114. Broman KW. R/qtlcharts: interactive graphics for quantitative trait locus mapping. Genetics. 2015;199(2):359–61.

115. Amezquita RA, Lun ATL, Becht E, Carey VJ, Carpp LN, Geistlinger L, et al. Orchestrating single-cell analysis with Bioconductor. Nat Methods. 2020;17(2):137–45.

116. Riemondy KA, Sheridan RM, Gillen A, Yu Y, Bennett CG, Hesselberth JR. valr: Reproducible genome interval analysis in R. F1000Res. 2017;6:1025.

117. Livak KJ, Schmittgen TD. Analysis of relative gene expression data using real-time quantitative PCR and the 2(-Delta Delta C(T)) Method. Methods. 2001;25(4):402–8.

118. Ogle DHW, P.; Dinno, A. FSA: Fisheries stock analysis. 2019.

119. Wickham H. Easily install and load the ‘tidyverse’. 2017.

120. Wickham H. Ggplot2: Elegant graphics for data analysis. 2016;Springer-Verlag New York.

121. Moser HL, Doe AP, Meier K, Garnier S, Laudier D, Akiyama H, et al. Genetic lineage tracing of targeted cell populations during enthesis healing. J Orthop Res. 2018;36(12):3275–84.

