## Supplemental figures for "Dynamic transcriptional and epigenetic changes define postnatal tendon growth"

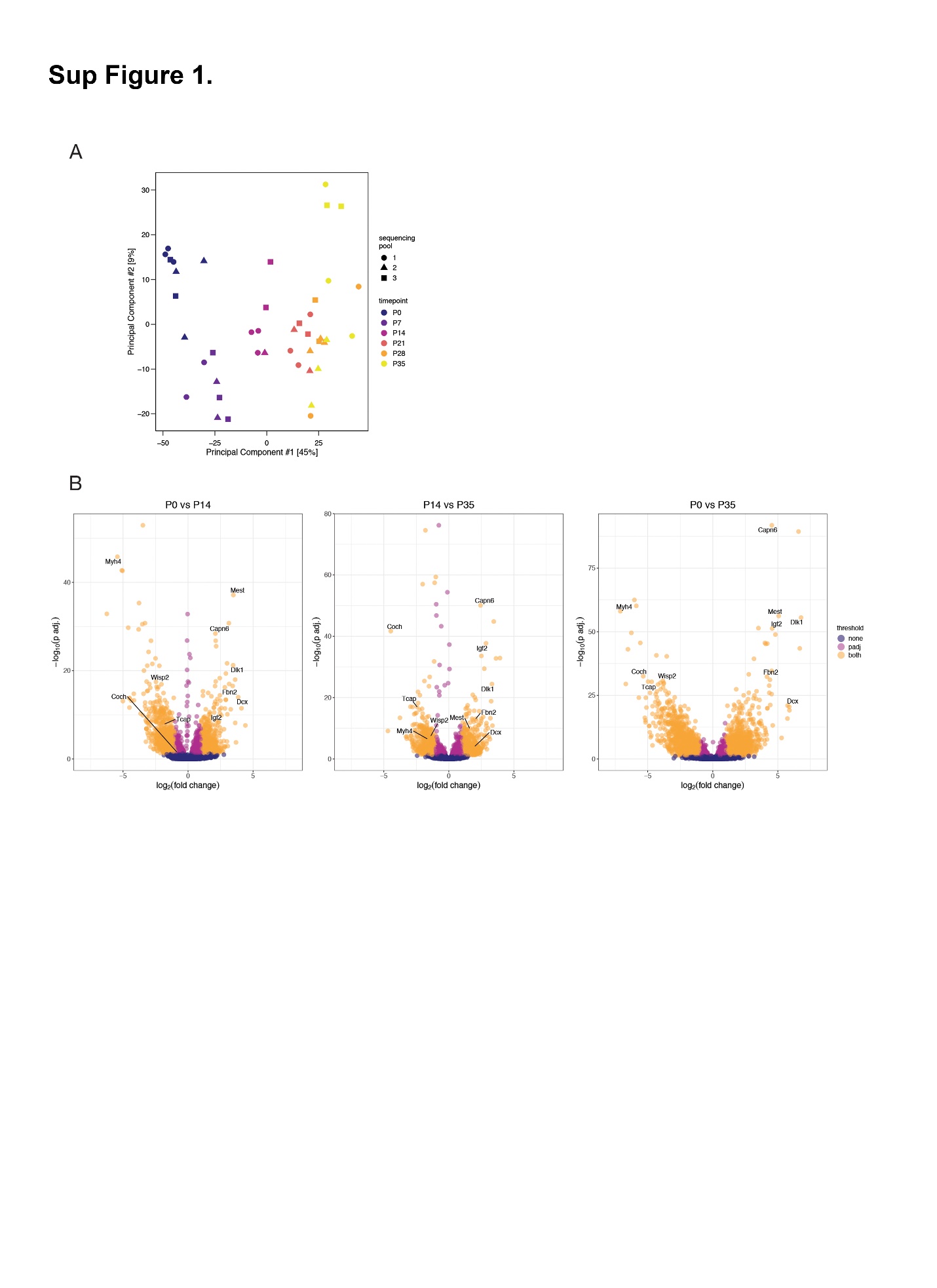
Supplemental Figures and Legends

Supplemental Figure 1

RNA-seq identifies differentially expressed genes at key time points during postnatal growth. A Principal Components Analysis (PCA) on normalized gene counts shows separation of samples along PC1 according to postnatal stage prior to P21 (A). Volcano plots showing differentially expressed genes between indicated timepoints (B).

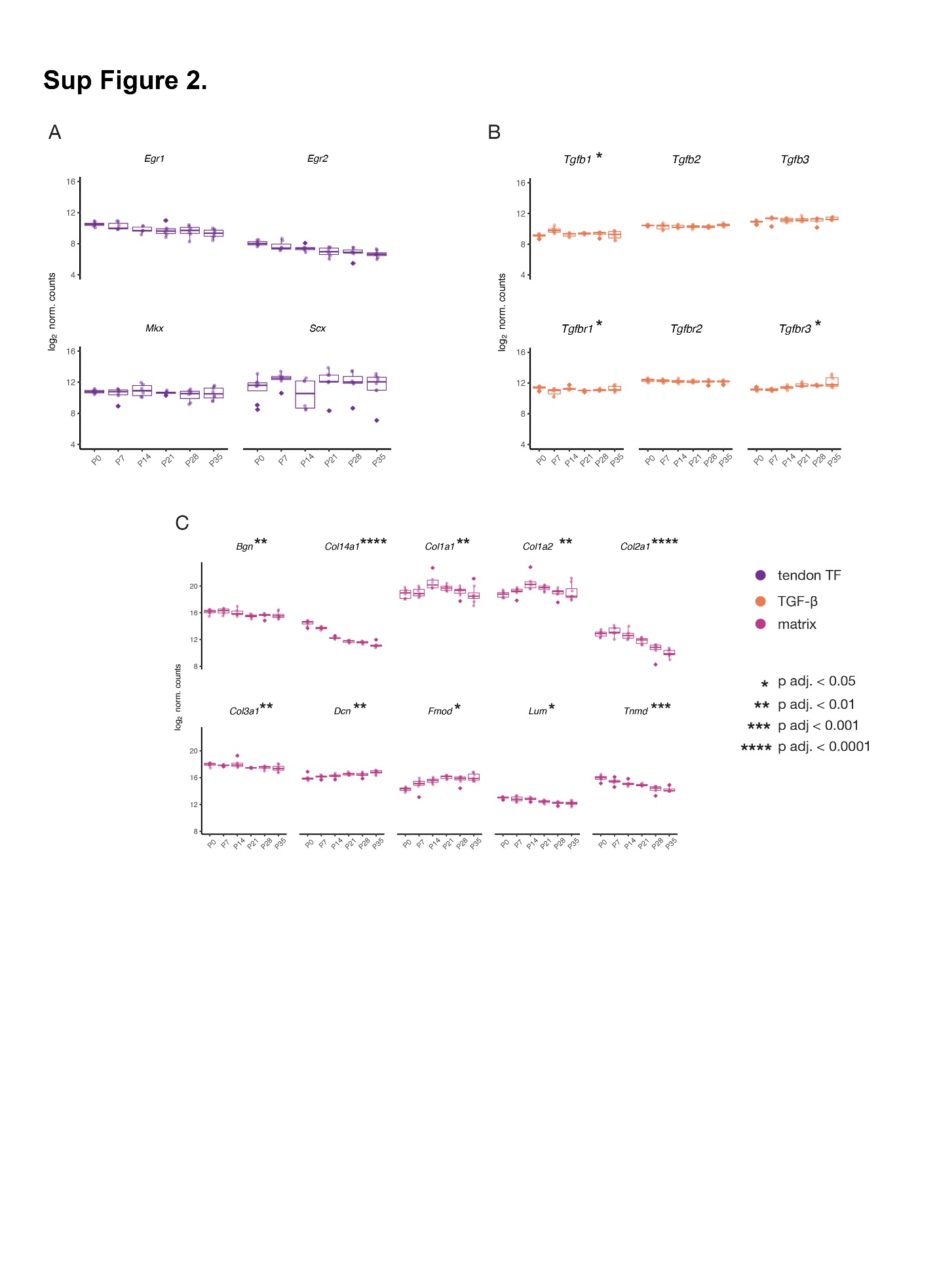
Supplemental Figure 2

Expression of known ‘tendon’ and ECM genes measured with RNA-seq. Genes associated with tendon development are not significantly differentially expressed at postnatal stages (A). Analysis of TGFβ ligands and receptors shows that *Tgfb1* is differentially upregulated at P7 only, *Tgfbr1* is intermittently differentially expressed, and *Tgfbr3* is differentially upregulated from P0 to P35 (B). Expression of ECM related genes are significantly differentially expressed in unique directions during postnatal stages with *Dcn* and *Fmod* increasing gradually over time, *Col2a1*, *Col14a1*, *Col3a1*, and *Tnmd* decreasing over time, *Col1a1* and *Col1a2* peak in expression at P14, and *Bgn* is expressed at higher levels from P00-P14 after which its levels decrease (C). Kruskal-Wallis rank sum test followed by a Dunn test with Benjamini-Hochberg correction were used to test for specific differences among pairs of time points.

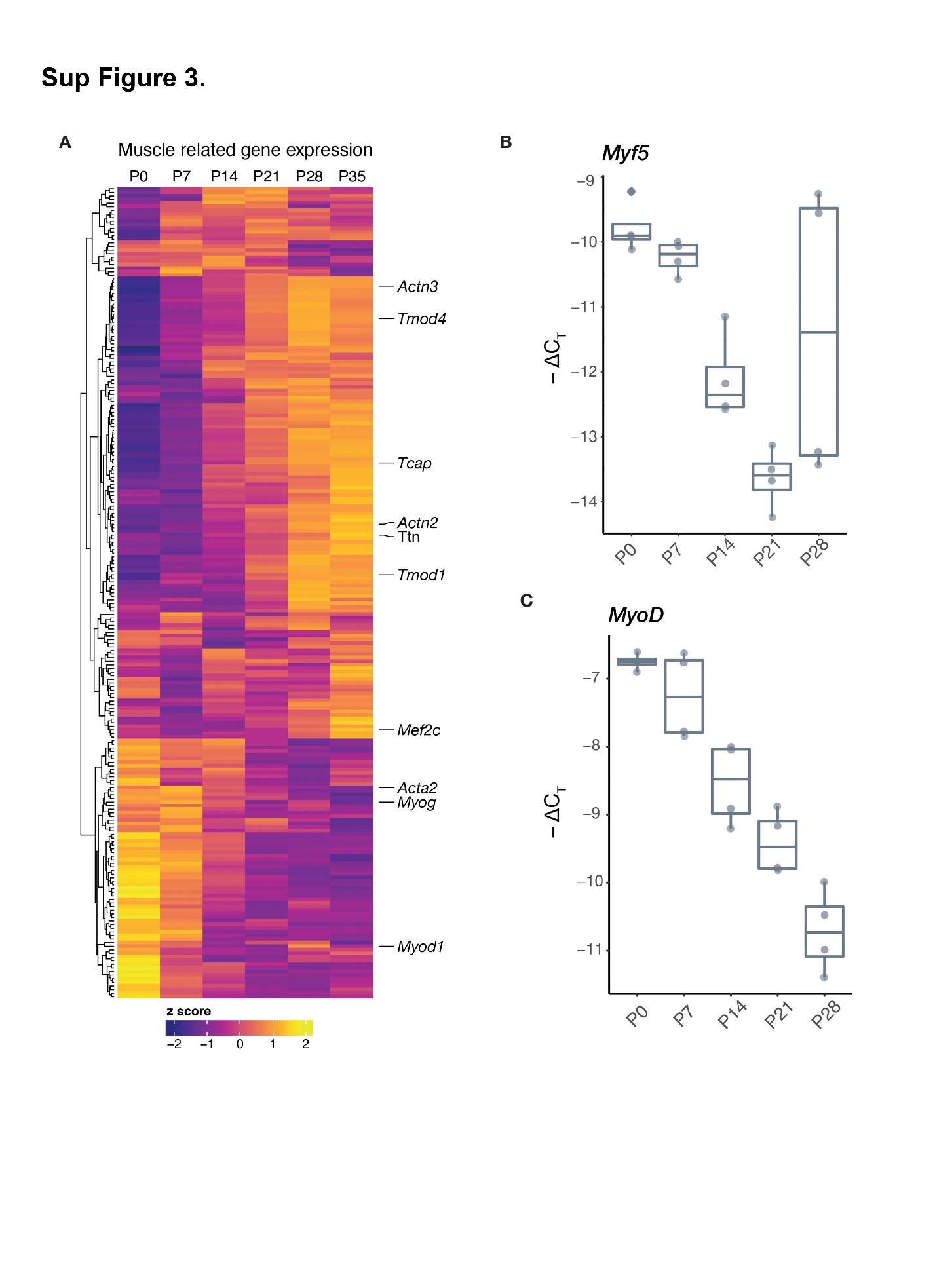
Supplemental Figure 3

Expression of ‘muscle’ genes in the tendon during early postnatal development. Heatmap showing RNA expression of muscle-associated genes in Clusters 1, 2, and 5 with specific genes indicated (A). RT-qPCR for *Myf5* and *MyoD* show decreased expression over postnatal time, validating the results of the RNA-seq (B). Kruskal-Wallis rank sum test followed by a Dunn test with Benjamini-Hochberg correction were used to test for specific differences among pairs of time points.

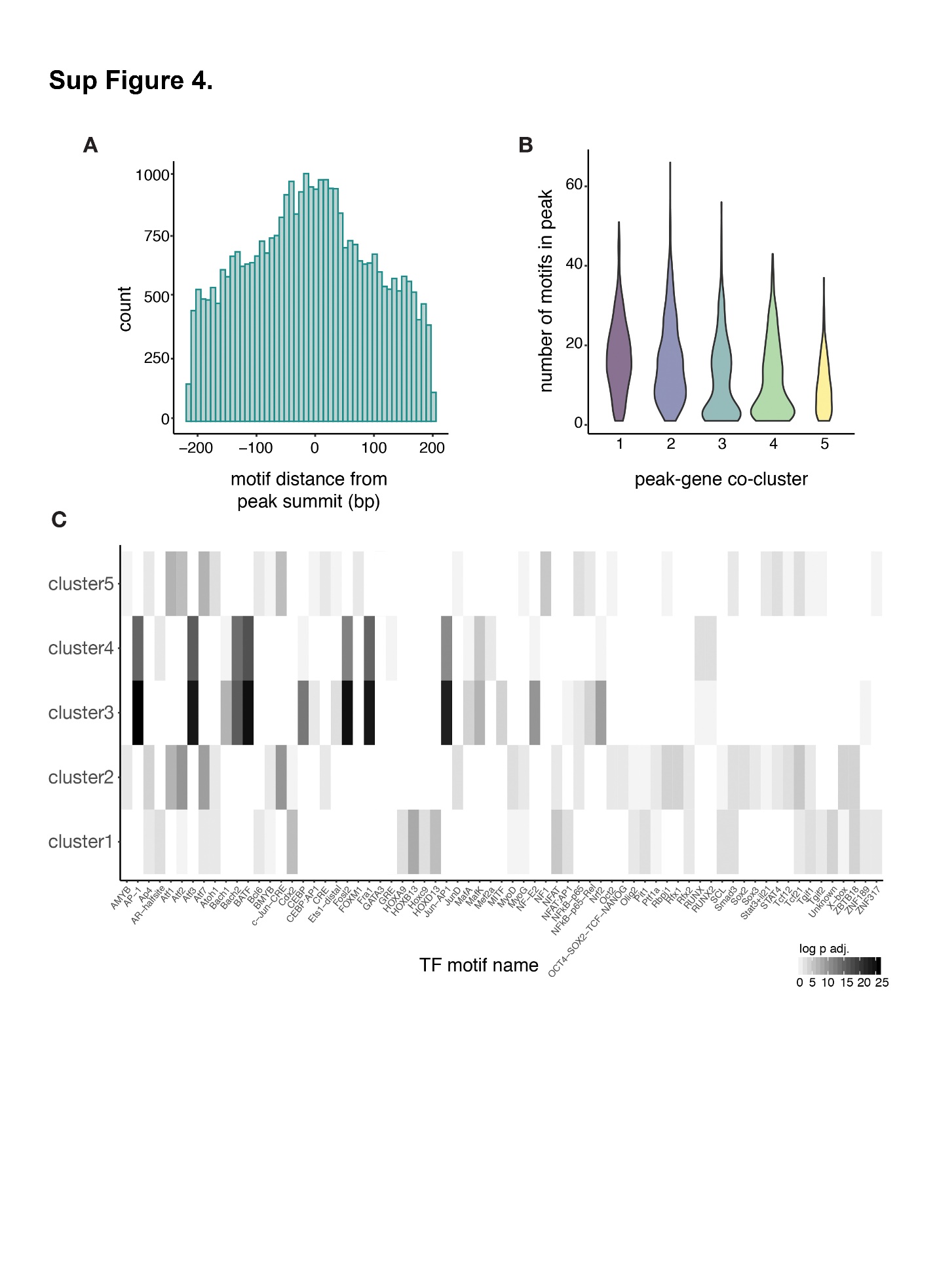
Supplemental Figure 4

Enrichment of transcription factor binding motifs within differentially accessible chromatin regions. The distribution of motifs relative to the peak summit shows that the majority of motifs are within 100 bp (A). The distribution in the number of motifs per peak shows only slight differences across clusters (B). Identification of motifs that are associated with specific clusters demonstrates shared motifs among clusters changing in the same direction with developmental timing (Clusters 1 and 2 or Clusters 3 and 4) but little overlap in those changing in opposite directions (Clusters 3 and 4 compared with Clusters 1, 2, and 5) (C).

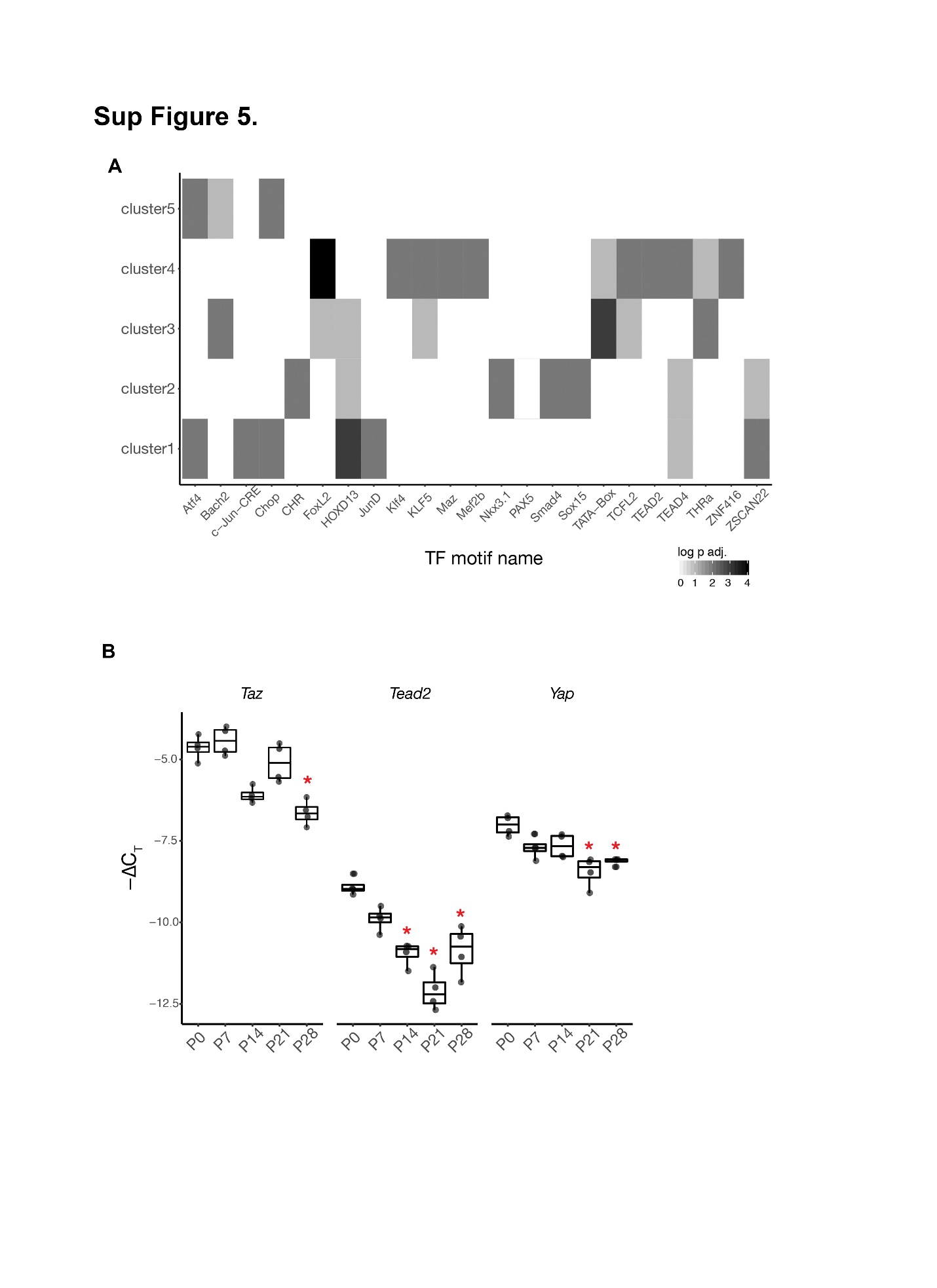
Supplemental Figure 5.

Enrichment of transcription factor binding motifs within the promoter regions of putative enhancer target genes (A). RT-qPCR analysis of Yap, Taz, and Tead2 shows a significant decrease in transcript levels over postnatal stages from P0 to P28. Statistical differences among the time points were investigated using a Kruskal-Wallis rank sum test followed by a Dunn test with Benjamini-Hochberg correction to test for specific differences among pairs of time points.

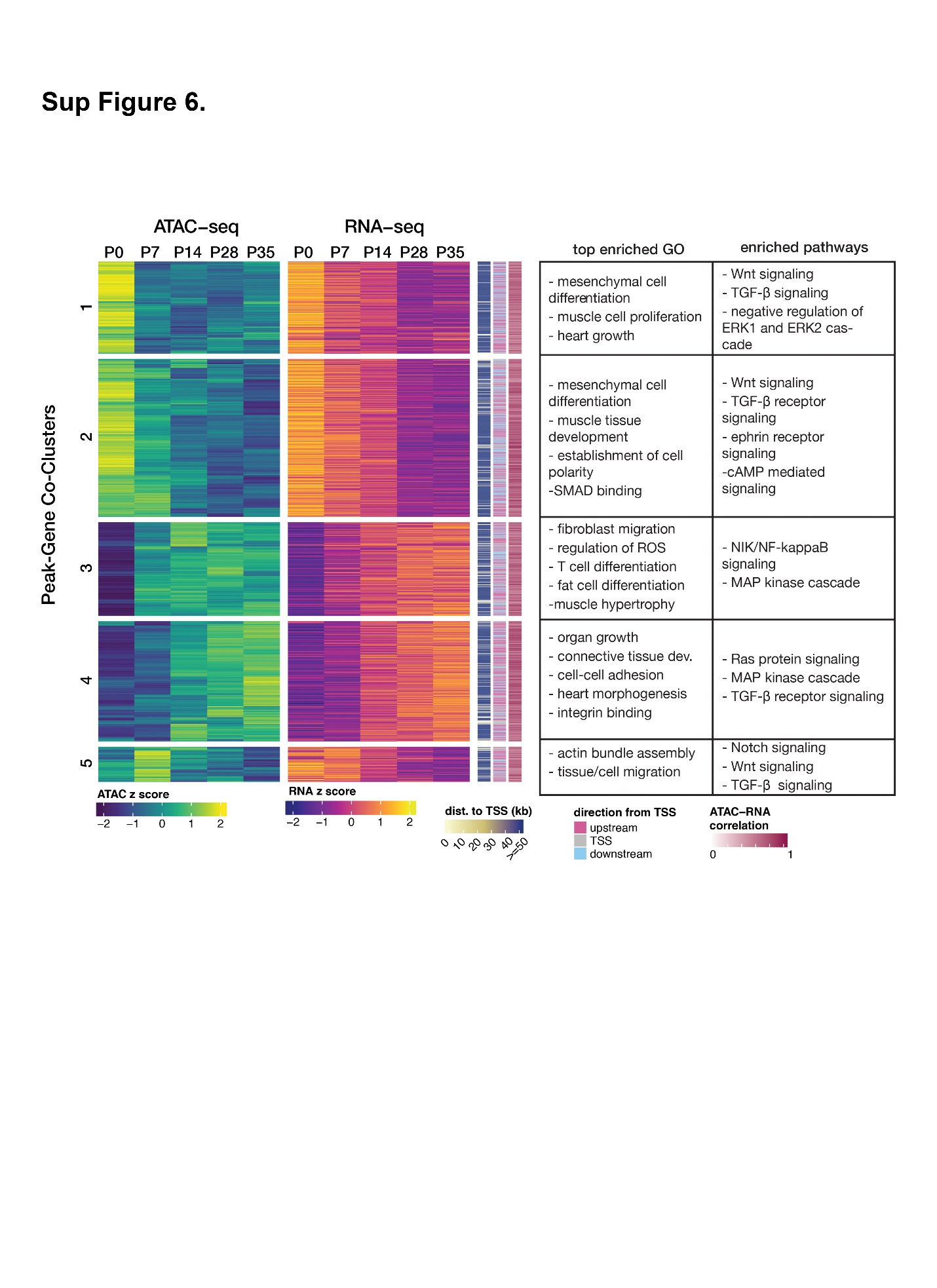
Supplemental Figure 6

Integrated RNA-seq/ATAC-seq analysis (same as in Figure 3) is shown with the top enriched GO terms and enriched pathways.

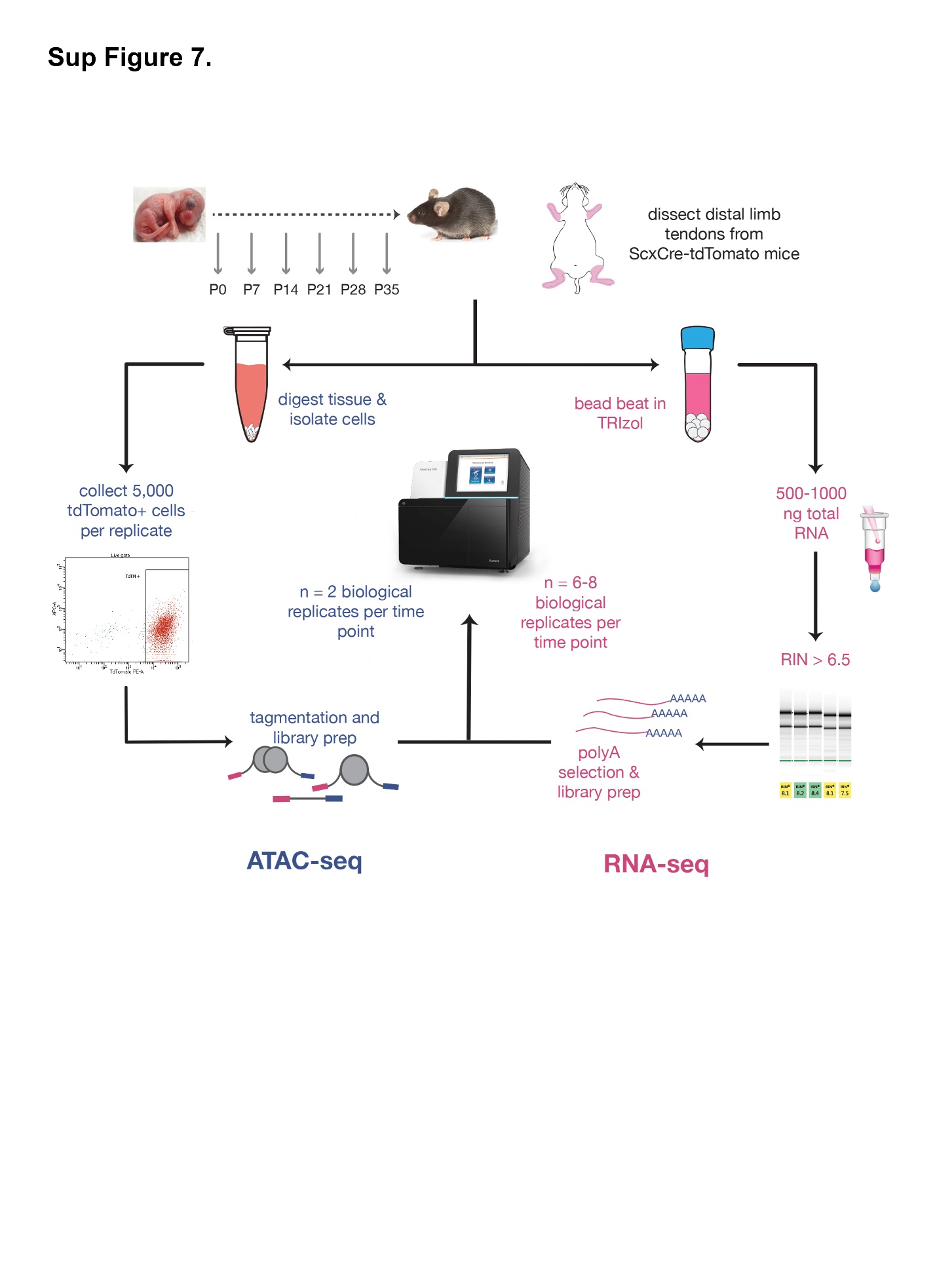
Supplemental Figure 7

Overview of method details for RNA-seq and ATAC-seq

**Supplementary Table 1:** **List of RT-qPCR primers**

| Target Gene | Forward Sequence (5’–3’) | Reverse Sequence (5’–3’) | Primer Source |
| --- | --- | --- | --- |
| *Col1a2* | CCAGCGAAGAACTCATACAGC | GGACACCCCTTCTACGTTGT | (Mendias et al., 2008) |
| *Gapdh* | TGTTCCTACCCCCAATGTGT | GGTCCTCAGTGTAGCCCAAG | (Szymaniak et al., 2015) |
| *Mki67* | AGCAAGCCAACAGAATTTCCAG | TATCTTGACCTTCCCCATCAGG | Self-designed using PrimerBlast |
| *Myf5* | CTGTCTGGTCCCGAAAGAAC | TGGAGAGAGGGAAGCTGTGT | (Shin et al., 2014) |
| *MyoD1* | TACAGTGGCGACTCAGATGC | GAGATGCGCTCCACTATGCT | (Hildyard and Wells, 2014) |
| *Taz* | GCCTGGCCTGCATTAAAATGG | CTTGCTTCAGAATTGGGCAGT | PrimerBank ID 21313658a1 |
| *Tead2* | GAGCCCCGACATTGAGCAG | CCGGCCATACATCTTGCCC | PrimerBank ID 7106433a1 |
| *Yap1* | AATGTGGACCTTGGCACACT | ACTCCACGTCCAAGATTTCG | (Szymaniak et al., 2015) |

**References:**

Hildyard, J.C., Wells, D.J., 2014. Identification and validation of quantitative PCR reference genes suitable for normalizing expression in normal and dystrophic cell culture models of myogenesis. PLoS Curr 6.

Mendias, C.L., Bakhurin, K.I., Faulkner, J.A., 2008. Tendons of myostatin-deficient mice are small, brittle, and hypocellular. Proc Natl Acad Sci U S A 105, 388-393.

Shin, S., Suh, Y., Zerby, H.N., Lee, K., 2014. Membrane-bound delta-like 1 homolog (Dlk1) promotes while soluble Dlk1 inhibits myogenesis in C2C12 cells. FEBS Lett 588, 1100-1108.

Szymaniak, A.D., Mahoney, J.E., Cardoso, W.V., Varelas, X., 2015. Crumbs3-Mediated Polarity Directs Airway Epithelial Cell Fate through the Hippo Pathway Effector Yap. Dev Cell 34, 283-296.
